# Engineered allostery in light-regulated LOV-Turbo enables precise spatiotemporal control of proximity labeling in living cells

**DOI:** 10.1101/2023.03.09.531939

**Authors:** Song-Yi Lee, Joleen S. Cheah, Boxuan Zhao, Charles Xu, Heegwang Roh, Christina K. Kim, Kelvin F. Cho, Namrata D. Udeshi, Steven A. Carr, Alice Y. Ting

## Abstract

The incorporation of light-responsive domains into engineered proteins has enabled control of protein localization, interactions, and function with light. We integrated optogenetic control into proximity labeling (PL), a cornerstone technique for high-resolution proteomic mapping of organelles and interactomes in living cells. Through structure-guided screening and directed evolution, we installed the light-sensitive LOV domain into the PL enzyme TurboID to rapidly and reversibly control its labeling activity with low-power blue light. “LOV-Turbo” works in multiple contexts and dramatically reduces background in biotin-rich environments such as neurons. We used LOV-Turbo for pulse-chase labeling to discover proteins that traffick between endoplasmic reticulum, nuclear, and mitochondrial compartments under cellular stress. We also showed that instead of external light, LOV-Turbo can be activated by BRET from luciferase, enabling interaction-dependent PL. Overall, LOV-Turbo increases the spatial and temporal precision of PL, expanding the scope of experimental questions that can be addressed with PL.

## Introduction

Many natural enzymes are controlled by allosteric regulation. Binding by a ligand, RNA, or protein at a distal site leads to conformational changes that are transmitted, sometimes over long distances, to the enzyme’s active site to alter its catalytic properties^1^. Among engineered enzymes, however, allostery is far less common^2–4^. Though allosteric control could improve the spatial and/or temporal precision of many engineered enzymes and enable input-dependent activity that could be exploited for synthetic circuit design, there are few systematic approaches for introducing engineered allostery into enzymes of interest.

Here we report the engineering of a light-regulated variant of TurboID via the introduction of designed allostery. TurboID is an artificial enzyme used for promiscuous proximity-dependent biotinylation in living cells and mapping of spatial proteomes^5^. Engineered via directed evolution from the *E. coli* enzyme biotin ligase (BirA), TurboID uses biotin and ATP to generate a reactive biotinyl-AMP mixed anhydride which it releases from its active site to covalently tag nearby proteins on nucleophilic lysine sidechains. While TurboID has been used extensively for mapping interactomes and organelle proteomes in culture and *in vivo*^6, 7^, it does have important limitations. First, TurboID activity is most commonly controlled by the availability of its substrate, biotin, which is endogenously present in most living organisms, contributing to background labeling prior to exogenous biotin addition and consequently, limited temporal specificity. Second, the spatial specificity of TurboID (and all other PL enzymes) is determined by the quality of its genetic targeting, which is sometimes insufficiently specific for the cell type, cellular subcompartment, or protein complex of interest.

To address these limitations, we report a light-regulated variant of TurboID, “LOV-Turbo”, a single 50 kD polypeptide that has nearly undetectable activity in the dark state but turns on within seconds of weak blue light illumination to give activity comparable to that of TurboID. We show that LOV-Turbo improves the spatial and temporal precision of proximity labeling in culture and *in vivo*, and enables new applications not possible with TurboID, including pulse-chase labeling to map proteome dynamics in living systems. Furthermore, our study demonstrates a semi-rational approach to engineering light-induced allostery with an extremely high dynamic range that could be applied to other enzymes of interest.

## Results

### Design of light-regulated TurboID, or LOV-Turbo

Light is a versatile input that can be delivered in a spatially and temporally precise manner, even *in vivo*. A light-regulated version of TurboID would enable precise control over the time window of proteome labeling as well as its location. We sought to introduce light regulation to TurboID with several design goals in mind. First, we envisioned a single protein construct, rather than a multicomponent system, for ease of use and robust performance across a range of expression levels. Second, to be maximally effective, light-regulated TurboID should have negligible “leak” or background activity in the dark state, and activity comparable to that of the parent TurboID enzyme in the presence of light. Third, light-regulated TurboID should be rapidly reversible, so that activity can be shut off after the desired labeling period, enabling “chase” experiments to be performed. Lastly, the illumination conditions required for TurboID activation should be mild (~1 mW/cm^2^), so that specialized equipment is not needed, and phototoxicity is minimal.

To meet these criteria, we envisioned integrating the LOV photosensory domain, a 16 kD flavin-containing protein from the *Avena sativa* oat plant, into TurboID to regulate its activity with light. The core of LOV forms a tight complex, or “clamp”, with its C-terminal Jα helix in the dark state but releases the Jα helix in the presence of blue 470 nm light (**Fig. 1A**). The light power required is extremely low; cell cultures containing the LOV domain can often be stimulated by ambient room light.

**Fig. 1.**
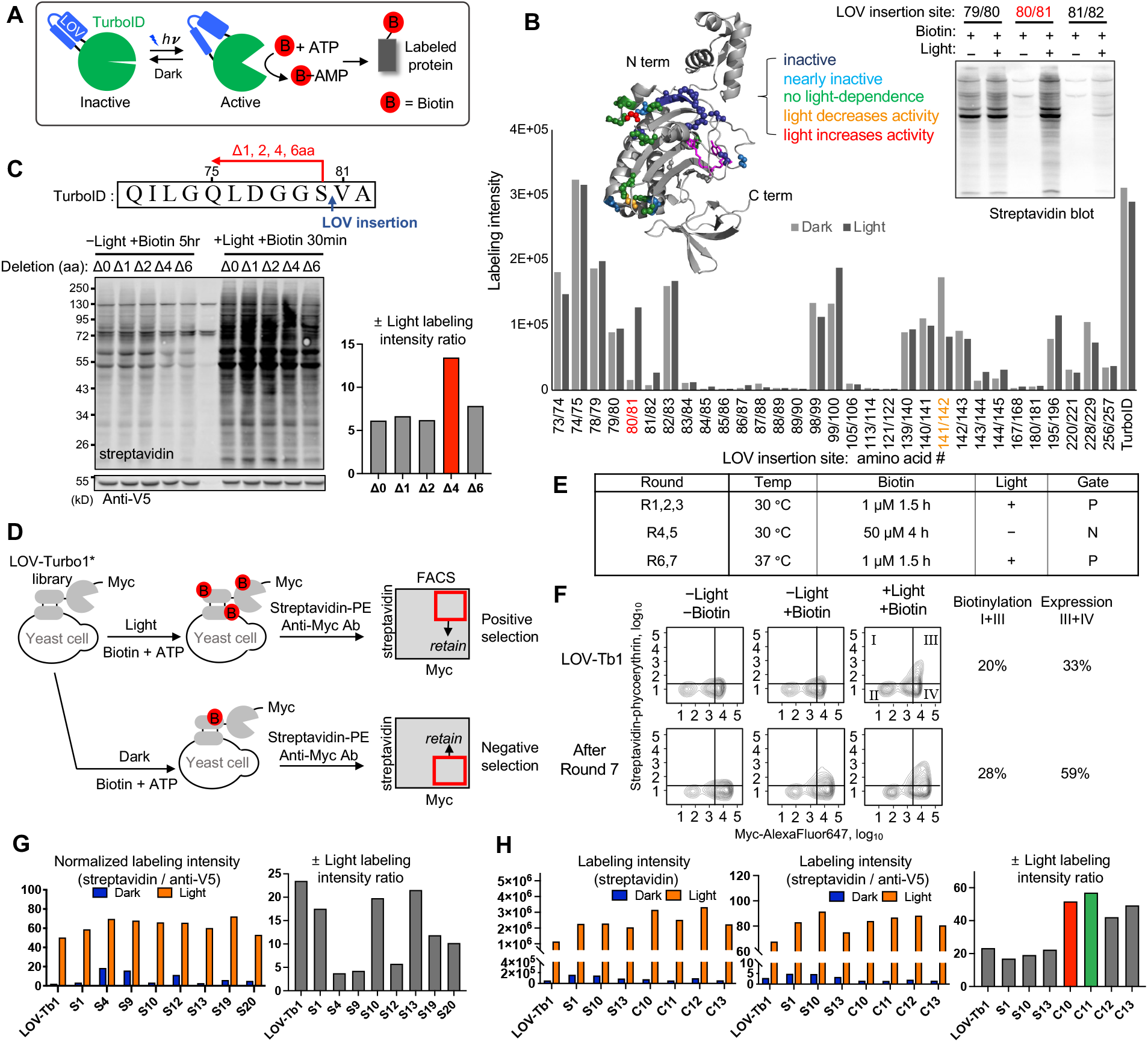
Design and directed evolution of LOV-Turbo. (**A**) LOV-Turbo has a blue-light sensitive hLOV1^9^ domain inserted into its 6th surface-exposed loop (domain structure shown in **Supplementary Fig. 1A**). In the dark state, clamping by hLOV1 keeps TurboID inactive, while in the light state, clamp release restores TurboID activity. Active LOV-Turbo promiscuously biotinylates nearby proteins, enabling their identification by mass spectrometry. (**B**) Screening of 31 LOV insertion sites in TurboID. Constructs were expressed in the HEK cytosol and labeling was performed for 30 minutes in light or dark. Relative biotinylation activity was quantified by streptavidin blotting of whole cell lysates. Inset shows sample data. Ribbon structure of BirA (PDBID: 2EWN) shows LOV insertion sites and the result obtained for each. Non-hydrolyzable biotinol-5’-AMP in pink. (**C**) Truncation of LOV-TurboID at the 80/81 insertion site improves light gating. Amino acids 75-80 were progressively truncated as shown. The Δ4 construct showed the highest ± light signal ratio and lowest -light leak. (**D**) Scheme for directed evolution on the yeast cell surface. PE, phycoerythrin. Ab, antibody. FACS, fluorescence-activated cell sorting. (**E**) Selection conditions used over 7 rounds. Positive selection (P) and negative selection (N) gates are shown in (D). (**F**) FACS plots comparing LOV-Turbo1 template to post-round 7 yeast population. Percentages give fractions of active cells (in quadrants I + III) and fractions of expressing cells (in quadrants III + IV). (**G**) Screening of 8 evolved mutants in the HEK cytosol. Biotinylation intensity quantified by streptavidin blot of whole cell lysates (shown in **Supplementary Fig. 1F**), after labeling for 30 minutes in light or 5 hours in dark. LOV-Tb1, LOV-Turbo1. (**H**) Same as (G) but with combination mutants C10-C13 as well, after labeling for 30 minutes in light or 4 hours in dark. Blots are shown in **Supplementary Fig. 1H**. C10 was selected over C11 as our final LOV-Turbo due to its higher expression level in HEK 293T cells.

By inserting LOV into a surface-exposed loop of TurboID that is allosterically coupled to its active site, we envisioned distorting and impairing the active site through LOV’s clamping effect in the dark state but releasing it to a “native-like” and active conformation upon illumination. To test this possibility, we selected 9 different surface-exposed loops in TurboID, based on the crystal structure of the parent enzyme BirA^8^, and explored 1-6 different LOV insertion sites in each loop (**Supplementary Fig. 1A**). A total of 31 insertion sites were tested, using the hybrid LOV domain (hLOV1^9^) that exhibits a combination of high signal in the light state and minimal leak in the dark state when used in other optogenetic tools such as FLiCRE^10^ and SPARK^9^.

12 out of 31 sites we tested eliminated TurboID activity altogether, and another 5 impaired activity in *both* light and dark states (**Fig. 1B** and **Supplementary Fig. 1A**). One site (141/142) showed light-dependent activity, but in the direction opposite to what we desired (activity inhibited by light). A single fusion construct showed increased activity after light exposure: the 80/81 LOV insertion, between the 4th α-helix and a mixed β-sheet of TurboID. In general, we found that loops closer to TurboID’s N-terminus were better able to tolerate LOV insertion than loops near the C-terminus. Also, insertion sites designed to block the biotin binding pocket resulted in a complete loss of TurboID activity.

We further optimized the 80/81 LOV-Turbo fusion to improve ± light signal ratio. The addition of Gly-Ser (GS) linkers N-terminal to the LOV domain had no effect, while GS linkers inserted C-terminal to LOV, after the Jα helix, were deleterious for light-dependence (**Supplementary Fig. 1B**). Even a single glycine inserted after Jα abolished the light-dependent activity of LOV-Turbo. Given this sensitivity, we hypothesized that it might be beneficial to further shorten the LOV insertion region. We did so by truncating the 80/81 loop into which the LOV domain was inserted, removing 1-6 amino acids N-terminal to Gly80. Removal of 4 amino acids decreased dark state activity while maintaining light state activity (**Fig. 1C**), giving more than a 2-fold improvement in ± light signal ratio.

This optimized construct, named LOV-Turbo1, was effective in cells, but we found that its expression level was considerably lower than that of TurboID, implying instability, and limited the levels of achievable biotinylation (**Supplementary Fig. 1C**). To address this problem, we turned to directed evolution, both to improve LOV-Turbo1’s expression and stability, and to further optimize its dynamic range.

### Directed evolution of LOV-Turbo

We selected yeast display as our directed evolution platform for LOV-Turbo, as it has previously been used to evolve TurboID^5^ and APEX2^11^. We further modified the platform to incorporate alternating rounds of positive selection in light and negative selection in the dark (**Fig. 1D**). By this approach, we hoped to enrich LOV-Turbo variants with a high dynamic range as well as robust cell surface expression.

We used error-prone PCR to introduce mutations throughout the LOV-Turbo1 template. Sequencing showed an average mutation rate of 1.4 amino acids per gene in our library. ~10^7^ LOV-Turbo variants were displayed on the yeast surface as a fusion to the Aga2p mating protein. We performed labeling with biotin and ATP in the presence of light for 90 minutes, before washing and staining with streptavidin-phycoerythrin (SA-PE) to detect self-biotinylation, and anti-Myc antibody staining to quantify LOV-Turbo expression level. Approximately 38% of our initial library had detectable biotinylation activity under these conditions, with a mean activity ~2-fold lower than that of LOV-Turbo1, indicating a good level of mutagenesis and library diversity.

We performed three rounds of positive selection, followed by two rounds of negative selection in the dark, enriching clones that had low streptavidin but high-Myc staining under these conditions. Lastly, we performed two more rounds of positive selection, but at 37 °C rather than the typical 30 °C yeast culturing conditions to increase selection stringency for stable and well-folded clones (**Fig. 1E** and **Supplementary Fig. 1D**). FACS plots in **Fig. 1F** show that our final round 7 yeast pool displays both higher expression and higher promiscuous biotinylation activity than the template, LOV-Turbo1. Sequencing of clones from this pool showed mutations in both the LOV domain and the TurboID gene, with T52S, I147T, I231V, and F286L mutations especially enriched (**Supplementary Table 1**). As shown in **Fig. 2A**, these mutations are distributed across the LOV-Turbo predicted structure, with some proximal to the biotin-binding site, and others in the N-terminal domain, far from the active site.

**Fig. 2.**
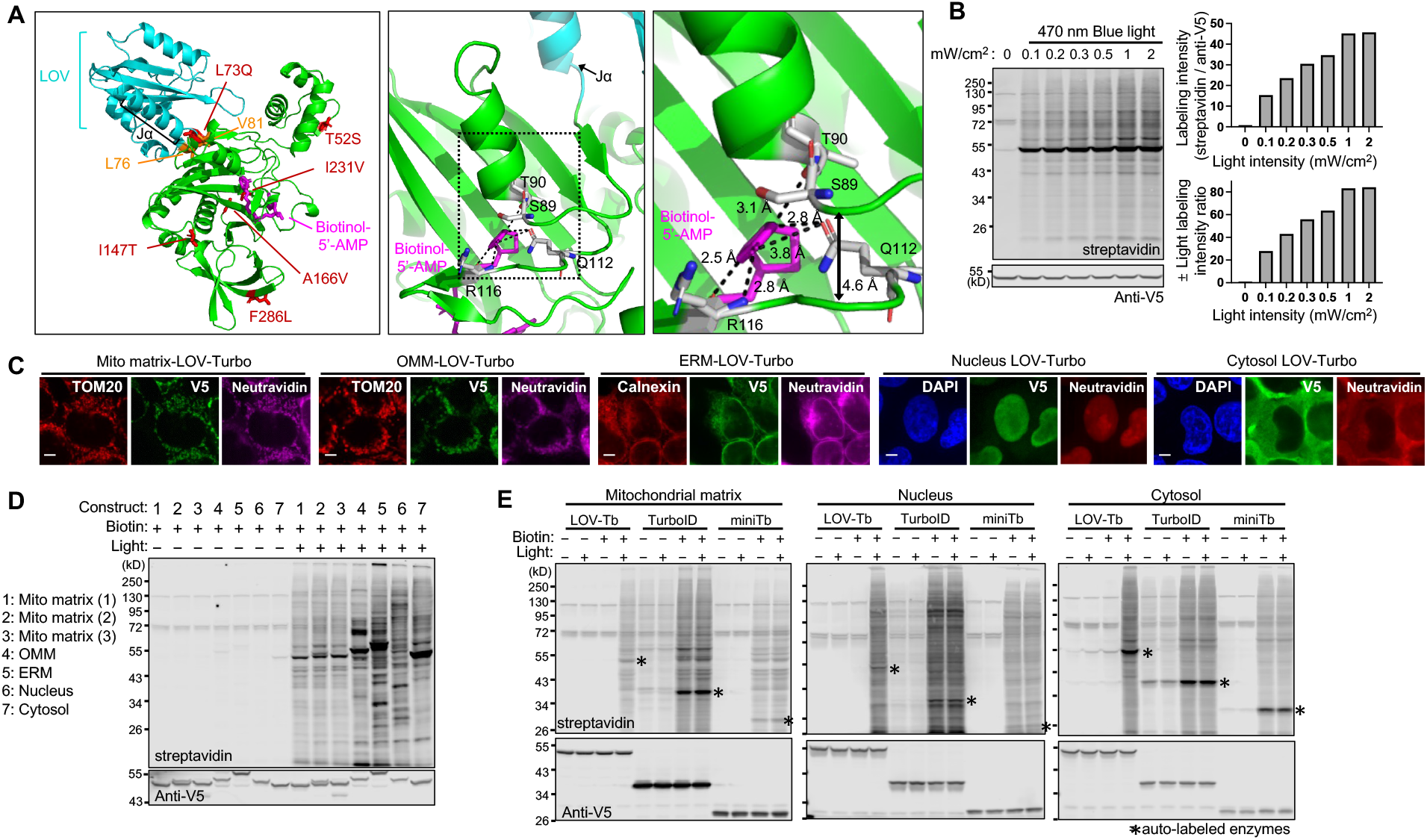
Characterization of LOV-Turbo. (**A**) AlphaFold-predicted structure of evolved LOV-Turbo (clone C10). LOV domain in cyan and TurboID in green. L76 and V81 flanking the LOV insertion site are colored orange. The 6 mutations enriched by directed evolution are colored red. Non-hydrolyzable biotinol-5’-AMP in pink. Close-up views show TurboID’s active site in relation to residues that may be allosterically coupled to the LOV domain. Double-headed arrow shows the distance between indicated backbone loops. (**B**) Light intensity-dependence of LOV-Turbo. HEK 293T cells expressing LOV-Turbo in the cytosol were stimulated with blue light of varying intensities (0.1 - 1 mW/cm^2^) for 15 minutes in the presence of biotin. Lysates were analyzed by streptavidin and anti-V5 blotting. (**C**) Confocal imaging of LOV-Turbo-catalyzed biotinylation in the mitochondrial matrix, outer mitochondrial membrane (OMM), ER membrane facing the cytosol (ERM), nucleus, and cytosol. Anti-V5 detects LOV-Turbo expression. Neutravidin-AF647 detects biotinylated proteins, after 30 minutes of labeling under blue light illumination. TOM20 and calnexin are endogenous mitochondrial and ER markers, respectively. DAPI stains the nucleus. Scale bars, 5 μm. (**D**) Streptavidin blotting of HEK 293T lysates expressing LOV-Turbo targeted to various compartments and labeled with biotin and blue light for 30 minutes. Three different mitochondrial matrix targeting sequences were tested (see **Methods**). (**E**) Comparison of LOV-Turbo (LOV-Tb), TurboID, and miniTurbo (miniTb) in HEK mitochondrial matrix, nucleus, and cytosol. Labeling time was 30 minutes. * indicates auto-labeled LOV-Turbo, TurboID or miniTurbo.

We characterized 19 clones from round 7 on the yeast surface (**Supplementary Fig. 1E**) and selected 8 for analysis in HEK 293T cells. All showed light-dependent activity in the HEK cytosol, but to varying degrees (**Fig. 1G** and **Supplementary Fig. 1F**). We used this data to guide the selection of mutations for manual recombination, generating another 13 LOV-Turbo variants (C1-C13, **Fig. 1H** and **Supplementary Fig. 1G-H**). Testing of these alongside yeast-enriched clones revealed that C10, with 6 mutations in the TurboID sequence and no mutations in the LOV domain, showed higher ±light signal ratio and expression level than LOV-Turbo1 template under matched conditions. The AlphaFold^12^-predicted structure of C10 (**Fig. 2A**) shows that its 6 mutations are distributed across the entire protein, with only two (I231V and A166V) close to the biotin/ATP substrate binding pockets. Other mutations may contribute to C10’s improved expression.

Using C10 as a template, we also tested if swapping out the entire hLOV1 domain for other published LOV domains could improve performance. **Supplementary Fig. 1I** shows that only iLID^13, 14^ and variants of hLOV (f-hLOV1^10^ (I25V or V14T)^15^ and hLOV1^9^), which are known to exhibit tight caging in the dark state, gave strong light gating. Wild-type AsLOV2 and eLOV^16^ constructs exhibited dark state leak. Ultimately, our original C10 with hLOV1 remained superior. We selected C10 as our final optimized LOV-Turbo, henceforth named “LOV-Turbo”.

### Characterization of LOV-Turbo in mammalian cells

To characterize LOV-Turbo, we performed a labeling time course in the HEK cytosol as well as a titration of biotin concentration. With 100 μM exogenous biotin, we observed a steady increase in promiscuous labeling over 120 minutes, with not much increase in signal beyond 150 minutes (**Supplementary Fig. 2A**). The labeling after 10 minutes was comparable to that of TurboID, and exceeded the signal from miniTurbo^5^, suggesting that the standard labeling time of 10 minutes used for TurboID should be suitable for LOV-Turbo as well (**Supplementary Fig. 2B**). We could hardly detect promiscuous labeling in the dark state, even at the highest biotin concentration of 0.5 mM, or when cells were incubated with 100 μM biotin overnight (**Supplementary Fig. 2C-D**). As expected, LOV-Turbo can be activated by 470 nm blue light, but not 660 nm red light, which facilitates the handling of LOV-Turbo samples in red-light-illuminated dark rooms (**Supplementary Fig. 2E**). We performed a study of LOV-Turbo light requirements and found the best turn-on with continuous illumination at 1 mW/cm^2^ power (**Fig. 2B**). Alternatively, 10-1000 ms pulses of 2.5 mW/cm^2^ light at a 33% duty cycle give a comparable degree of activation (**Supplementary Fig. 2F-G**).

We next probed LOV-Turbo performance across multiple subcellular compartments in HEK 293T cells (**Fig. 2C-E**). We observed strong light- and biotin-dependent labeling in the cytosol, nucleus, mitochondrial matrix, outer mitochondrial membrane, and endoplasmic reticulum membrane facing the cytosol. Each compartment gave a unique “fingerprint” by streptavidin blotting, reflective of the different subcellular proteomes tagged by LOV-Turbo in each location (**Fig. 2D**). In the cytosol, LOV-Turbo was more active than TurboID, whereas TurboID activity was higher in the nucleus and mitochondrial matrix (**Fig. 2E**).

Interestingly, we found that LOV-Turbo was much less active in the ER lumen and on the surface of HEK 293T cells (**Supplementary Fig. 2H-J**), which contrasts with its activity on the yeast cell surface, where we performed our directed evolution. We speculated that the oxidative environment of the ER lumen could potentially lead to intra- or intermolecular disulfide formation that disrupts TurboID folding and/or activity. We therefore tested mutation of Cys103 in TurboID or Cys128 in the LOV domain; however, neither Cys removals restored activity in the ER (**Supplementary Fig. 2J**).

### LOV-Turbo is reversible and works in multiple cell types

To test if light-activated LOV-Turbo can reverse to the inactive state when light is removed, we compared biotinylation activity by LOV-Turbo when light is supplied for 15 minutes and then turned off, versus when light is continually on for 1 hour. If LOV-Turbo turns on but then does not reverse when the light is removed, then labeling will continue in the presence of biotin even after the 15-minute light window. **Fig. 3A** shows that this does not occur. We further probed the kinetics of LOV-Turbo turn-on and turn-off in HEK 293T cells (**Supplementary Fig. 3A-B**). To study turn-on, during a 5-minute light stimulation window, we supplied biotin at the 0, 1, 2, or 3-minute mark. The biotinylation intensity progressively decreased, indicating that LOV-Turbo turns on within a minute of light application, consistent with the reported LOV kon of 2 ms^17^ (**Supplementary Fig. 3A**). To probe turn-off kinetics, we provided light, followed by biotin 0 or 1 minute later (**Supplementary Fig. 3B**). Our results indicate that LOV-Turbo is reversible, and turn-on and turn-off are both fast, occurring within seconds of light application and light removal.

**Fig. 3.**
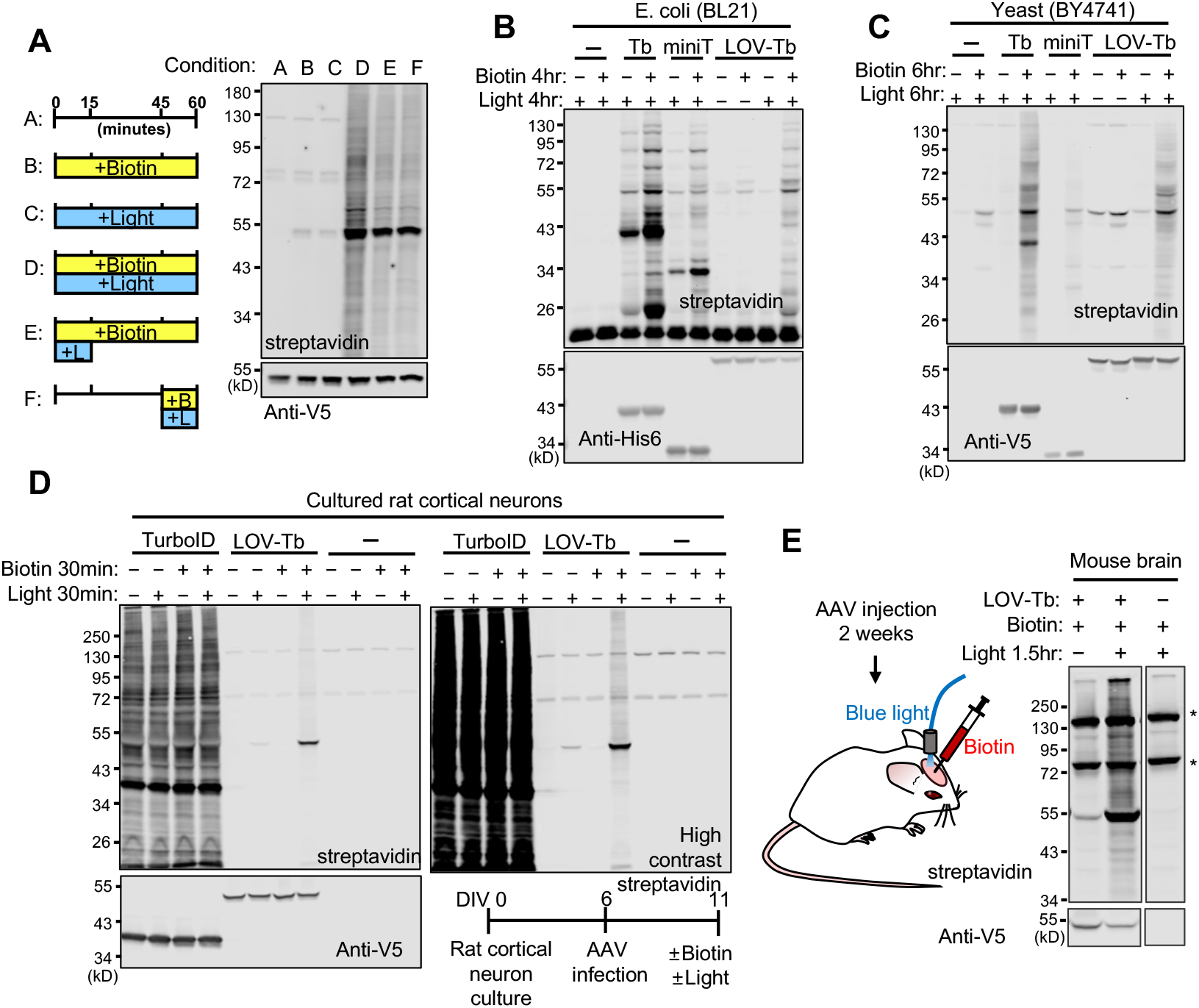
LOV-Turbo is reversible and works in multiple cell types and in the rodent brain. (**A**) LOV-Turbo labeling is terminated by the removal of light. HEK 293T cells expressing LOV-Turbo in the cytosol were labeled as shown in conditions A-F. Whole-cell lysates were then analyzed by streptavidin blotting. (**B**) LOV-Turbo labeling in *E. coli.* Constructs were expressed in the cytosol of BL21 *E. coli* and labeling was performed for 4 hours. Anti-His6 antibody detects ligase expression. Tb, TurboID. miniT, miniTurbo. (**C**) LOV-Turbo labeling in yeast (BY4741 strain). Labeling was performed for 6 hours. (**D**) LOV-Turbo labeling in cultured rat cortical neurons. LOV-Turbo or TurboID was expressed in the cytosol via adeno-associated virus 1/2 (AAV) infection for 5 days prior to labeling. TurboID shows high background even without exogenous biotin addition. DIV, days *in vitro.* (**E**) LOV-Turbo labeling in the mouse brain. Cytosolic LOV-Turbo was expressed in the mouse cortex via AAV1/2 injection. 2 weeks later, 0.5 μL of 10 mM biotin was injected into the brain while 470 nm light (5 mW/cm^2^, 10 ms pulses at 10Hz) was delivered for 1.5 hours. Whole-tissue lysate analyzed by streptavidin and anti-V5 blotting. * indicates endogenously biotinylated proteins.

Finally, we tested if LOV-Turbo can be used for light-dependent proximity labeling in other cell types and *in vivo* (**Fig. 3B-E**). We expressed LOV-Turbo in the cytosol of either bacterial or yeast cells, and performed 4 and 6 hours of labeling, respectively, in the presence of blue light and 100 μM biotin. Promiscuous biotinylation was observed in both contexts, with activity comparable to TurboID and miniTurbo, while negligible labeling was observed in omit-light controls (**Fig. 3B-C**). LOV-Turbo was also active in cultured rat cortical neurons, although a fair comparison to TurboID could not be performed due to the high level of biotin in the culture media, which made it impossible to control the TurboID labeling time window (**Fig. 3D** and **Supplementary Fig. 3C**). Lastly, we tested LOV-Turbo activity in the intact mouse brain (**Fig. 3E**). Adeno-associated virus containing the LOV-Turbo gene was injected into the primary motor cortex of adult mice, and 2 weeks later, we delivered blue 470 nm light via an implanted optical fiber for 1.5 hours while biotin was simultaneously injected into the brain. Promiscuous labeling of endogenous proteins was observed, but not in the omit-light controls. These studies illustrate the versatility of LOV-Turbo across multiple organelles and cell types.

### Applications enabled by LOV-Turbo

LOV-Turbo improves the temporal specificity of proximity labeling. In many settings, TurboID is able to use the low levels of biotin present in culture media or *in vivo* to initiate labeling prior to the addition of exogenous biotin. By contrast, LOV-Turbo gives negligible labeling, even in the presence of 0.5 mM biotin, if the light is not supplied (**Supplementary Fig. 2D**). This difference is particularly pronounced in biotin-rich neuron culture, where TurboID samples show the same level of streptavidin or neutravidin staining whether or not exogenous biotin is added (**Fig. 3D** and **Supplementary Fig. 3C**). Furthermore, when targeted to the mitochondria, the ongoing biotinylation activity of TurboID leads to auto-labeling of a critical lysine residue in the targeting sequence, causing mislocalization of the construct to the nucleus and catalysis of background biotinylation of nuclear proteins (**Fig. 4A**). By contrast, LOV-Turbo in the same context is correctly targeted and produces mitochondria-specific biotinylation. The improvement in mitochondria-specific biotinylation can be observed in HEK 293T cell cultures as well (**Supplementary Fig. 3D**), although some mistargeting is seen at high expression levels, perhaps due to the larger size and tightly folded structure of LOV-Turbo.

**Fig. 4.**
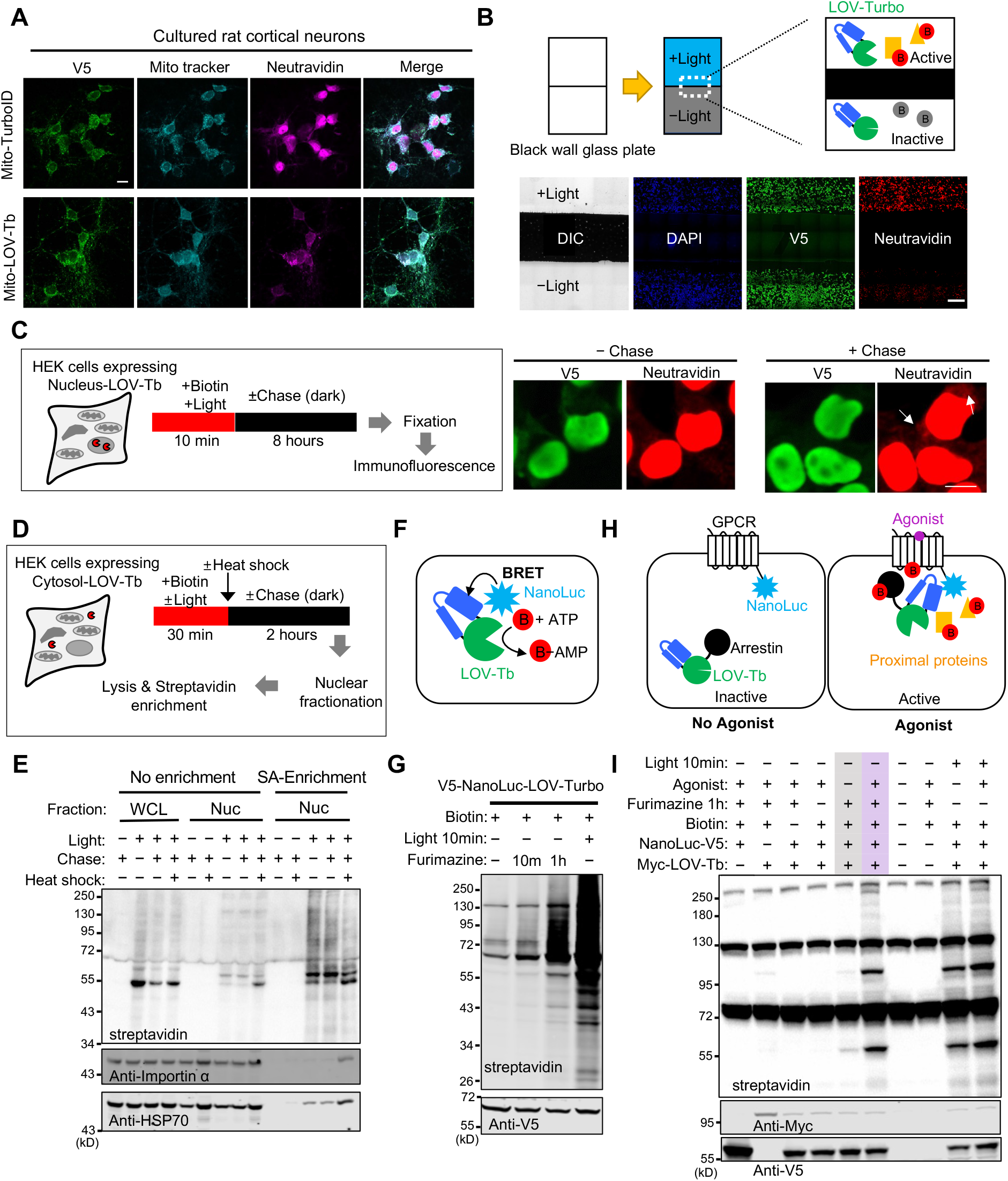
Applications of LOV-Turbo. (**A**) Reduced background and improved spatial precision with mito-LOV-Turbo in cultured rat cortical neurons. LOV-Turbo or TurboID targeted to the mitochondrial matrix were expressed in cultured rat cortical neurons via AAV1/2 transduction at DIV6. At DIV12, neurons were treated with biotin and light for 2 hours, then fixed and stained with anti-V5 antibody to detect enzyme expression, and neutravidin-AF647 to detect biotinylated proteins. Scale bar, 10 μm. (**B**) Spatial control of proximity labeling with LOV-Turbo. HEK 293T cells expressing cytosolic LOV-Turbo were labeled with biotin and light for 5 minutes, while half of the sample was covered with black tape. Images show neutravidin staining in lit area only. Scale bar, 500 μm. (**C**) Pulse-chase labeling with LOV-Turbo. HEK 293T cells expressing nuclear LOV-Turbo were labeled with biotin and light for 10 minutes, then chased for 8 hours in the dark (other chase times shown in **Supplementary Fig. 3G**). Images show neutravidin staining before and after the chase period. Arrows point to biotinylated proteins in the cytosol. Scale bar, 10 μm. (**D**) Pulse-chase labeling with cytosolic LOV-Turbo following heat shock. After a 2-hour chase at 37 °C or 42 °C, biotinylated proteins were enriched from purified nuclei. (**E**) Blots showing importin-α and HSP70 from the cytosol detected in the nucleus following heat shock. WCL, whole cell lysate. Nuc, nuclear fraction. SA, streptavidin. (**F**) Schematic of BRET-based activation of LOV-Turbo with the luciferase NanoLuc. (**G**) Testing a direct fusion of NanoLuc to LOV-Turbo. Addition of NanoLuc’s substrate, furimazine, in the dark is sufficient to activate LOV-Turbo and produce promiscuous biotinylation in the cytosol of HEK 293T cells. (**H**) Schematic of LOV-Turbo activation via arrestin recruitment to activated GPCR-NanoLuc fusion. (**I**) Specific proximity labeling by a GPCR-arrestin complex, via BRET-induced LOV-Turbo activation. Arrestin-LOV-Turbo and CCR6-NanoLuc were expressed in HEK 293T cells, and the GPCR (CCR6) was stimulated by its peptide agonist CCL20. Labeling was performed for 1 hour in the presence of biotin and furimazine. Anti-Myc detects arrestin-LOV-Turbo expression, and anti-V5 detects CCR6-NanoLuc expression.

To demonstrate spatial control of PL with LOV-Turbo, we illuminated a cell culture dish that was partially covered with black tape. Streptavidin staining was observed exclusively in the non-covered, illuminated regions of the sample (**Fig. 4B**).

A third application enabled by LOV-Turbo is pulse-chase labeling. TurboID labeling can be inhibited by cooling samples to 4 °C^5^, but if cells are maintained at 30 °C or 37 °C, biotin washout is not sufficient to terminate labeling (**Supplementary Fig. 3E**). Because LOV-Turbo is turned on *reversibly* by light (**Fig. 3A**), the mere removal of light terminates biotinylation. Cells can then continue to be cultured, as the LOV-Turbo-labeled proteome redistributes. We demonstrated this in **Fig. 4C** by performing 10 minutes of biotinylation with LOV-Turbo, then chasing for 8 hours in the dark. A small but noticeable population of biotinylated proteins can be observed in the cytosol after chasing for 8 hours but not immediately following biotinylation (**Fig. 4C** and **Supplementary Fig. 3F-G**).

Pulse-chase labeling with LOV-Turbo can be combined with organelle fractionation to query biotinylated proteins that have been redistributed into a specific organelle. We demonstrated this by labeling the cytosolic proteome of HEK 293T cells with LOV-Turbo-NES, heat shocking the cells, chasing for 2 hours in the dark, then performing nuclear fractionation and streptavidin (SA) enrichment to isolate biotinylated proteins in the nucleus (**Fig. 4D-E**). **Fig. 4E** shows a range of proteins captured by this method, including importin-α and HSP70, which are both known to translocate from the cytosol to the nucleus upon heat shock^18, 19^.

The fourth application of LOV-Turbo we explored is inspired by our previous observation that LOV-based optogenetic tools can be controlled by bioluminescence from luciferases via BRET^20^ (**Fig. 4F**). We first tested whether LOV-Turbo could be activated by BRET rather than exogenous blue light by directly fusing the 460 nm-emitting luciferase NanoLuc^21^ to LOV-Turbo. We observed biotinylation by this construct in HEK 293T cells when NanoLuc’s substrate, furimazine, was supplied together with biotin for 10 minutes or 1 hour in the dark (**Fig. 4G**). Next, we tested an intermolecular BRET configuration by fusing NanoLuc to the GPCR CCR6 and LOV-Turbo to arrestin, which is recruited to CCR6 upon activation by CCR6’s peptide agonist CCL2 0^22^ (**Fig. 4H**). **Fig. 4I** shows promiscuous biotinylation by arrestin-LOV-Turbo in the presence of furimazine and biotin, in cells treated with the CCL20 agonist but not in untreated cells. Controls omitting furimazine or biotin showed a lack of labeling. These results suggest that it may be possible to use LOV-Turbo and NanoLuc in combination to label the interactomes of defined protein subcomplexes in living cells.

### Mapping proteome dynamics with LOV-Turbo by mass spectrometry

We further explored the use of LOV-Turbo for pulse-chase labeling in living cells by performing two quantitative liquid chromatography-mass spectrometry (LC-MS/MS)-based proteomics experiments (**Fig. 5A** and **5C**). In both, we first used LOV-Turbo to tag the endogenous ER membrane-associated proteome from the cytosolic face. We then performed either nuclear fractionation or mitochondrial fractionation, after a chase period of several hours. In both experiments, we analyzed basal trafficking in addition to trafficking following ER stress. Treatment with either tunicamycin^23^ (for ER-to-nucleus) or thapsigargin^24^ (for ER-to-mitochondria) induced ER stress by inhibiting N-linked glycosylation or blocking sarco/ER Ca^2+^-ATPase, respectively.

**Fig. 5.**
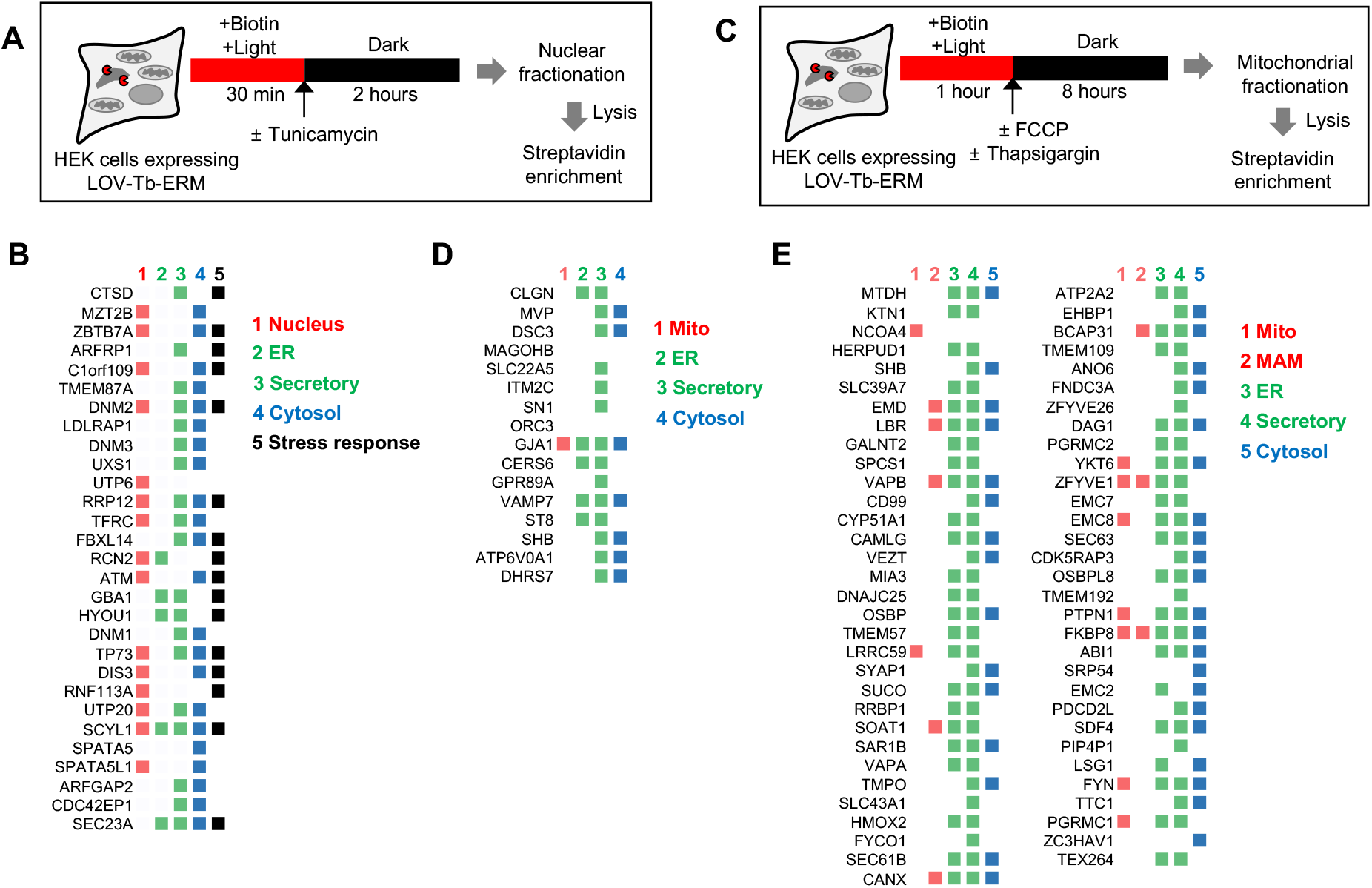
Mapping proteome dynamics with LOV-Turbo by mass spectrometry. (**A**) Scheme for pulse-chase labeling with LOV-Turbo localized to the ER membrane (ERM) facing the cytosol. After a 2-hour chase, biotinylated proteins were enriched from purified nuclei. See **Supplementary Fig. 4C** for design of 18-plex proteomic experiment. (**B**) 29 proteins that exhibit increased trafficking from ERM to nucleus following stress induction by tunicamycin. Proteins are ranked by fold-change compared to basal (no stress) condition. Colorings based on Gene Ontology Cell Component (GOCC) annotation (details in **Supplementary Table 2**). (**C**) Scheme for pulse-chase labeling with LOV-Turbo at the ER membrane (ERM) facing cytosol. After an 8-hour chase, biotinylated proteins are enriched from purified mitochondria. See **Supplementary Fig. 5B** for design of 17-plex proteomic experiment. (**D**) 16 proteins detected in ERM to mitochondria dataset from (C) under basal conditions. Proteins ranked by fold-change compared to FCCP control, which inhibits mitochondrial protein import. Coloring reflects GOCC annotations (details in **Supplementary Table 3**). (**E**) 63 proteins that exhibit increased trafficking from ERM to mitochondria following thapsigargin treatment. Proteins ranked by fold-change compared to basal condition. Colorings based on GOCC annotation (details in **Supplementary Table 3**). MAM, mitochondria-associated membranes, which includes mitochondria-ER contact site proteins.

We first used Western blotting to check the quality of our organelle-fractionated, streptavidin-enriched material. In our ER-to-nucleus sample, we detected ATF6, a known stress-responsive ER-localized transcription factor^25^ (**Supplementary Fig. 4A**). In agreement with literature^25, 26^, we found that nuclear ATF6 is cleaved after tunicamycin-induced ER stress. **Supplementary Fig. 5A** shows the population of proteins detected in our ER-to-mitochondria sample. Encouragingly, treatment of cells with FCCP, which depolarizes mitochondria and prevents protein import, reduced the quantity of proteins remaining after mitochondrial fractionation and streptavidin enrichment.

The design of our ER-to-nucleus proteomic experiment is shown in **Supplementary Fig. 4C**. Negative controls omitting light or the 2-hour chase, spatial references with cytosolic LOV-Turbo, and whole cell lysate were included in the 18-plex TMT labeling experiment. A total of 5308 proteins were detected and quantified LC-MS/MS with good correlation observed between replicates (**Supplementary Fig. 4D**). To check the specificity of the data, we plotted enrichment ratios (experiment vs. omit light, experiment vs. cytosolic reference, etc.) as receiver operating characteristic (ROC) curves, using known ER or nuclear proteins as true positives and known mitochondrial proteins as false positives (**Supplementary Fig. 4E-G**). We observed that ER and nuclear proteins were strongly enriched over mitochondrial proteins with high TMT ratios, helping to confirm the specificity of our dataset.

As shown in **Supplementary Fig. 4H**, the 5308 detected proteins were filtered by p-value and fold-change with respect to omit-light, omit-tunicamycin, cytosolic LOV-Turbo, and whole cell lysate samples, to obtain a final list of 29 proteins that show increased ER-to-nucleus trafficking after tunicamycin treatment (**Supplementary table 2**). Within this list, 20 proteins have secretory pathway annotation, 15 proteins have nuclear annotation, and 7 have both nuclear and secretory annotation (**Fig. 5B**). 16 out of 29 proteins are known stress-responsive proteins. Three of our hits have previously been reported to translocate to the nucleus under stress (Serine-protein kinase ATM^27^, Cathepsin D CTSD^28^ and Hypoxia up-regulated protein 1 HYOU1^29^). Especially, ATM is an ER stress-responsive protein^30^, though the localization is still controversial^31^. Our dataset also may contain potential artifacts, such as nuclear proteins that are translated by ER-associated ribosomes during the biotinylation time window (e.g., nucleolar protein UTP6).

Nevertheless, we were intrigued by N-terminal kinase-like protein SCYL1. This protein has both ER/Golgi annotation^32^ and nuclear annotation, as one of its isoforms may bind to the human TERT promoter^33^. Reticulocalbin-2 RCN2 also appears in our dataset. It is ER-annotated by Gene Ontology Cellular Component (GOCC)^34, 35^, and enriched in our previous ERM-APEX2 dataset^36^. Though its function is unknown, previous studies have shown that RCN2 levels decrease after thapsigargin treatment in Huh7 cells^37^. See **Supplementary Text 1** for further analysis of this dataset.

In our second proteomic experiment, we combined LOV-Turbo labeling at the ER membrane with mitochondrial fractionation. The design of the 17-plex proteomic experiment is shown in **Supplementary Fig. 5B**, incorporating controls omitting light or inhibiting mitochondrial protein import with FCCP. 5107 proteins were detected by LC-MS/MS with good correlation between replicates (**Supplementary Fig. 5C**), and we performed filtering as shown in **Supplementary Fig. 5E**. The ROC curves in **Supplementary Fig. 5D** show strong enrichment of ER-annotated true positive proteins over false positive nuclear proteins, in both basal and thapsigargin-treated datasets. After filtering, we obtained lists of 16 and 64 proteins that traffick from ER to mitochondria under basal and thapsigargin stress conditions, respectively. (**Supplementary table 3**)

Both lists are enriched in ER and secretory pathway-annotated proteins (**Fig. 5D-E**). We identified a number of proteins associated with mitochondria-ER contact sites, as our filtering procedure does not eliminate such proteins; these include EMD^38^, LBR^38^, VAPB^38^, SOAT1^38^, BCAP31^38^, FKBP8^38^, CANX^39^, and ZFYVE1^39^. We also observed some artifacts likely resulting from contamination of mitochondrial fractions with ER-derived vesicles (e.g., the ER transmembrane proteins MTDH, KTN1, HERPUD1). We were intrigued by the tyrosine-protein phosphatase PTPN1, found in our thapsigargin dataset. Annotated as both a peripheral ER membrane protein and a mitochondrial matrix protein by GOCC, PTPN1 is activated under ER stress and serves as a negative regulator for insulin and leptin signaling^40, 41^. It also modulates oxidative phosphorylation, and its inhibition has been shown to promote mitochondrial fusion^42^. If PTPN1 shuttles from the ER membrane into mitochondria upon stress, as suggested by our LOV-Turbo pulse-chase labeling, this could serve as a mechanism to coordinate the ER stress response with mitochondrial oxidative phosphorylation and/or mitochondrial dynamics.

## Discussion

Through a combination of structure-guided design, screening, and directed evolution, we have developed a reversible blue light-gated variant of the proximity labeling enzyme TurboID. LOV-Turbo is 50 kD, ~16 kD larger than TurboID (34 kD) and boasts an extremely wide dynamic range: activity is virtually undetectable in the dark state, even in the presence of 0.5 mM biotin (50-100 μM biotin is typically used for labeling), and light-state activity is comparable to TurboID. In this study, fold turn-ons of LOV-Turbo ranged from 50 to 170-fold, depending on labeling time and organelle. Light-induced turn-on is rapid and can be rapidly reversed as well by simply removing the light source. We showed that LOV-Turbo works in multiple mammalian subcompartments, in yeast and bacteria, and also in the mouse brain.

A common strategy for introducing light gating to engineered proteins is to split the protein into fragments, and attach the fragments to light-inducible dimerizing proteins such as CRY/CIBN^43^ or FKF1/GI^44^. Such an approach has been used to produce light-gated versions of TALEs (transcription activator-like effectors)^45^, botulinum toxin^46^, and Gal4^47^. However, such tools are more cumbersome to use, as two separate components must be introduced, and tool performance varies depending on expression ratio as well as absolute expression level – with low reconstitution/signal at low expression levels, and possible false positives, or light-independent background, at high expression levels. Though we have previously developed split-TurboID^38^, we opted not to use it as the basis of light-gated TurboID. Instead, we aimed for a single, compact construct that would be simpler to use and work robustly across a range of expression levels.

A number of studies have engineered allosteric regulation by small molecules or light into proteins of interest, most commonly transcription factors^45, 48^. There are fewer examples of engineered allosteric regulation of enzymes^2–4^. In our own previous work^3^, we introduced calcium-binding domains into a distal surface-exposed loop of TEV protease to produce a Ca^+2^-activated protease that was used in a transcriptional Ca^+2^ integrator tool called scFLARE. Here we show that this approach is more broadly applicable. Some screening is required to identify a sensitive site for the insertion of protein domains that are conformationally responsive to analytes or light. Examination of the AlphaFold-predicted structure of LOV-Turbo (**Fig. 2A**) suggests that the 80/81 loop works because it is connected by a ß-strand to the biotin-binding pocket of TurboID. The “clamped” form of the hLOV1 domain in the dark state may distort TurboID’s biotin-binding pocket just enough to prevent substrate binding, while light releases the clamp and restores TurboID’s structure to a native-like conformation. Future structural and kinetic analyses of LOV-Turbo may reveal the structural features and enzymatic sub-steps that are most influenced by light.

A recent report described photo-Turbo, a different light-gated version of TurboID^49^. With a nitrobenzyl-caged catalytic lysine, introduced by unnatural amino acid mutagenesis, photo-Turbo is much more complicated to use, not genetically encoded, and not reversible. Uncaging also requires 365 nm UV light which can be phototoxic. More recently, a few reports have described light-catalyzed proximity labeling using mutants of the LOV domain (LOV*)^50, 51^. High power (>30 mW/cm^2^) blue light is used to induce singlet oxygen generation by LOV*, which in turn produces biotin-phenoxyl radicals for labeling. Our LOV-Turbo requires far weaker light for activation (~1 mW/cm^2^), no special media, and labeling is performed with biotin which is easier to deliver to cells, tissues, and animals than biotin-phenol. The low power requirement of LOV-Turbo enables the luciferase/BRET-based turn-on shown in **Fig. 4F-I**, which we do not believe would be possible with LOV*.

We have demonstrated several applications of LOV-Turbo, capitalizing on the improved temporal and spatial control afforded by light gating. LOV-Turbo suppressed background labeling prior to exogenous biotin addition that can plague TurboID experiments, especially *in vivo.* We utilized the reversible light regulation of LOV-Turbo to perform pulse-chase proteomics, identifying proteins that may shuttle between the ER and nucleus or mitochondria under normal and stressed conditions. Finally, we showed that instead of external blue light, LOV-Turbo can be activated by the luciferase NanoLuc via BRET, enabling us to selectively biotinylate protein subcomplexes, defined by proximity between a LOV-Turbo-fusion protein and a NanoLuc-fused protein.

It is also important to note LOV-Turbo’s limitations. Unlike TurboID, LOV-Turbo is inactive in the secretory compartment of mammalian cells. It also displays inferior targeting to some compartments such as the mitochondrial matrix, perhaps a result of its larger size and the high stability of the hLOV domain. Finally, further engineering will be required to improve the efficiency of BRET-based activation of LOV-Turbo by NanoLuc, before this approach can be widely applied for subcomplex-specific proteome mapping.

## Acknowledgements

Rat cortical neurons were a kind gift from Michael Lin (Stanford University). We thank Dr. Jochen Reinstein (Max Planck Institute) for helpful feedback. This work was supported by the NIH (R01-DK121409 and RC2DK129964 to A.Y.T, R01-OD026223 to A.Y.T. and S.A.C, and T32GM007276 to J.S.C.), the Stanford Wu Tsai Neurosciences Institute (A.Y.T.), the NSF (NeuroNex grant 2014862 to A. Y. T. and GRFP DGE-1656518 to J.S.C.), the National Research Foundation of Korea Grant NRF-2019R1A6A3A03033677 (S.-Y.L.), the Stanford Gerald J. Lieberman Fellowship (J.S.C.), and the Burroughs Wellcome Fund CASI 1019469 (C.K.K.). A.Y.T. is an investigator of the Chan Zuckerberg Biohub.

## Methods and Materials

### Labeling in mammalian cells with LOV-Turbo

HEK 293T cells transiently or stably expressing LOV-Turbo or its variants under a pTRE3G promoter were induced with 100-200 ng/ml doxycycline overnight. After induction, cells were covered with aluminum foil until ready for blue light stimulation. When uncovered, cells were handled in a dark room with red light illumination to avoid undesired activation of LOV-Turbo. To stimulate LOV-Turbo, biotin was added to cell cultures to a final concentration of 100 μM (unless noted otherwise) and incubated 37 °C while placed directly on top of a blue light LED array. We used an AMUZA system consisting of a blue LED array, an LED Array Driver and pulse generator. We used a 10% duty cycle, 10 ms on/ 90 ms off, for 30 minutes unless noted otherwise. Light power before pulse generation was typically ~10 mW/cm^2^ (measured by Thorlabs PM100D), unless noted otherwise. As shown in **Fig. 2B**, continuous blue light at 1 mW/cm^2^ activates LOV-Turbo just as well. Alternatively, 10 ms to 1 s pulses of 2.5 mW/cm^2^ light at a 33% duty cycle give a comparable degree of activation, as shown in **Supplementary Fig. 2G**. Note that we used 12-well cell culture plates (Corning, #3513) to test required light power, which may vary in thickness across different plate manufacturers. Control samples were processed in parallel omitting light or biotin. Following light stimulation, cells were once again handled under red light, washed in ice-cold DPBS three times, and analyzed by Western blot or immunofluorescence as described below.

### Western blot detection of LOV-Turbo activity

Following light stimulation, cells were washed in cold DPBS three times and lysed directly in the cell culture wells with RIPA lysis buffer supplemented with 1x protease inhibitor cocktail (Sigma-Aldrich). Lysates were cleared via centrifugation at 20,000 rcf at 4 °C for 10 minutes. Cleared lysates were mixed with protein loading buffer and boiled at 95 °C for 5 minutes. Samples were then loaded on 10% SDS-PAGE gels, transferred onto nitrocellulose membrane, and stained with Ponceau S (Sigma) to visualize total protein loading. Blots were blocked in 5% (w/v) nonfat milk (Lab scientific, M-0841) in 1x TBST (Teknova) for 30-60 minutes at room temperature, incubated with mouse anti-V5 primary antibody against LOV-Turbo’s epitope tag in TBST for 1 hour at room temperature, washed 3 times with TBST for 5 minutes each, incubated in anti-mouse-IRDye680 secondary antibody and streptavidin-IRDye800 or streptavidin-HRP for 30 minutes at room temperature, washed 3 times with TBST, and imaged on an Odyssey CLx imager (LI-COR) or ChemiDoc XRS+ imager with Clarity Western ECL Blotting Substrates (Bio-Rad). See antibody sources and dilutions used in Table “Antibodies used in this study”. Quantification of labeling (in **Figures 1B**, **1C**, **1G**, **1H**, and **2B**, and **Supplementary Figures 1G**, **1I**, **2C**, **2D**, **2F**, **2G**, and **3A**) was performed with Image Studio (LICOR) except for **Fig. 1B** which used Image J (NIH), by selecting the entire gel lane and subtracting endogenous biotinylated protein bands.

### Immunofluorescence detection of LOV-Turbo activity

Prior to labeling, cells were plated on glass coverslips coated with 25 μg/mL human fibronectin in DPBS for 1 hour at 37°C. After LOV-Turbo labeling, cells were fixed with 4% (v/v) paraformaldehyde in DPBS for 15 minutes, washed three times with DPBS, and permeabilized with ice-cold methanol at 4 °C for 10 minutes or with 0.1% Triton X-100 in DPBS at room temperature for 10 minutes. Cells were then washed three times with DPBS and blocked for 1 hour with 2% BSA (w/v) in DPBS at room temperature. Samples were incubated with primary antibodies in 1% BSA in TBST for 1-2 hours at room temperature. We used anti-V5 antibody to detect LOV-Turbo’s epitope tag, antibodies against TOM20 and calnexin to visualize mitochondria and ER, respectively, and other antibodies as listed in Table “Antibodies used in this study”. Sometimes, we used Mitotracker (Invitrogen, M22426) to visualize mitochondria following the manufacturer’s protocol in live cells. After primary antibody incubation, cells were washed three times in TBST and then incubated with secondary antibodies and neutravidin conjugated to AlexaFluor-488/568/647 in 1% BSA in TBST for 1 hour at room temperature. Cells were washed three times with TBST and then incubated with 300 nM DAPI in DPBS for 5 minutes. Cells were washed three times with DPBS and mounted on glass slides and imaged by confocal fluorescence microscopy (Zeiss). Images were collected with Slidebook (Intelligent Imaging Innovations).

### Mammalian cell culture, transfection, and generation of stable cell lines

HEK 293T cells (from ATCC) were cultured in DMEM containing 10% fetal bovine serum, 100 U/mL penicillin, and streptomycin at 37 °C and 5% CO2. For transient expression (**Fig. 1B-C** and **4G**, and **Supplementary Fig. 1B, 1F-H** and **2H-J**), cells in 12 well plates were typically transfected at ~70% confluency using 2.5 μL of polyethyleneimine (PEI, 1 mg/mL in water, pH 7.3) or Lipofectamine 2000 (Life Technologies) and total 1000 ng of plasmid in 100 μL serum-free media. Cells were incubated with 1 mL complete media supplemented with 100-200 ng/mL doxycycline overnight. Stable cell lines were generated with lentivirus infection. Lentivirus was generated by transfecting HEK 293T cells plated to ~60% confluency in a 6-well dish with 250 ng of pMD2G, 750 ng psPAX2, and 1000 ng lentiviral vector containing the gene of interest with 10 μL polyethyleneimine (1 mg/mL in water, pH 7.3) in serum-free media. Lentivirus-containing supernatant was collected after 48 hours and filtered through a 0.45 μm filter. HEK 293T cells at ~60% confluency were infected with crude lentivirus followed by selection with 10 μg/mL puromycin for a week. To induce expression of proteins under doxycycline-inducible promoter, cells were incubated with media supplemented with 100-200 ng/mL doxycycline overnight.

### Analysis of LOV-Turbo activity on the yeast surface

Yeast cells expressing surface-displayed LOV-Turbo as Aga2 fusions in a pCTCON2^52^ vector were cultured overnight in biotin-depleted media (BDM, see Yeast cell culture section) to induce expression. 6E9 cells (1 OD600 = 1E7 cells) were transferred into 1 mL of BDM containing 1 mM ATP, 5 mM MgCl2, and 1-50 μM of biotin. Cultures were then incubated at 30°C or 37°C for 1.5 hours with or without light exposure, while rotating. To keep samples in the dark, tubes were wrapped in aluminum foil. For light stimulation, samples were exposed to ambient light plus an additional blue light bulb overhead. After biotin labeling, samples were kept on ice to stop the labeling. Yeast cells were washed in cold PBS containing 0.1% BSA (PBS-B) three times by pelleting at 3,000 rcf for 2 minutes at 4 °C and resuspending in 1 mL cold PBS-B. Yeast cells were stained in 50 μL of PBS-B containing anti-Myc antibody (1:400) for 1 hour at 4 °C for detecting ligase expression, then the cells were washed three more times and stained in 50 μL PBS-B with goat anti-chicken-AF488 or anti-chicken-AF647 and streptavidin-phycoerythrin (SA-PE) for 40 min at 4 °C. Samples were washed for the final three times with 1 mL PBS-B before analysis or sorting. To analyze and sort, single yeast cells were gated on a forward-scatter area (FSC-A) by side-scatter area (SSC-A) plot around the clustered population (P1) on a ZE5 Cell Analyzer (Bio-Rad). P1 was then gated on a side-scatter width (SSC-W) by side-scatter height (SSC-H) plot around the clustered population (P2). P2 populated cells were then plotted to detect AF488 or AF647 and PE signals. Quantification of biotinylation in **Fig. 1F and Supplementary Fig. 1E** was calculated as the percent of yeasts in P2 in the upper quadrants I and III as drawn in **Fig. 1F**. Quantification of expression **Fig. 1F and Supplementary Fig. 1E** was calculated as the percent of yeasts in P2 in the right-most quadrants III and IV.

### Yeast display library generation

The LOV-Turbo mutant library was generated using error-prone PCR with 100 ng of LOV-Turbo1 as the template. Two libraries were generated at different levels of mutagenesis under 2 μM 8-oxo-dGTP and either 2 or 1 μM dPTP with 15 PCR cycles following published protocols^53^ with the following primers:

F: 5’-ggaggctctggtggaggcggtagcggaggcggagggtcggctagc-3’
R: 5’-gatctcgagctattacaagtcctcttcagaaataagcttttgttcggatcc-3’

PCR products were gel purified then reamplified for 30 more cycles under normal conditions and gel purified again. The pCTCON2 backbone was digested with NheI and BamHI and gel purified as well. Both 4000 ng of PCR product and 1000ng of cut vector were mixed and dried in a DNA speed vacuum (DNA110 Savant). The dried DNA was then resuspended in 10 μl of water and electroporated into electrocompetent EBY100 yeasts. After electroporation, yeasts were rescued in 2 mL of yeast extract peptone dextrose media (YPD) and recovered at 30 °C without shaking for 1 hour. 1.98 ml of the culture was then propagated in 100 mL of SDCAA while 20 μL was used to determine library size. Yeasts were diluted 100x, 1000x, 10000x, 100000x and 20 μl of each dilution was plated onto SDCAA plates at 30 °C for 3 days. The resulting library size was determined by the number of colonies on the 100x, 100x, 1000x, 100000x plates, corresponding to 10^4^, 10^5^, 10^6^, or 10^7^ transformants in the library, respectively. The library sizes resulting from the 2 levels of mutagenic error-prone PCR were 3.4E7 and 2.6E7. Both libraries were combined in equal volume to form our yeast display library.

### Directed evolution of LOV-Turbo on yeast

The two LOV-Turbo libraries generated above were combined 1:1. For the first round of selection, yeast cells were labeled with 1 μM of biotin for 1.5 hours at 30 °C then subsequently processed by staining with primary and secondary antibodies as described above. To sort on a BD FACS Aria II cell sorter (BD Biosciences), gates for singlets were drawn on an FSC-A by SSC-A plot to generate population 1 (P1). P1 was then gated on an FSC-A and FSC-H to generate population 2 (P2), and then P2 was gated on an SSC-A and SSC-W plot to further filter for singlets to generate population 3 (P3). P3 was then gated on a AF647 by SA-PE plot to collect high expressing and high activity clones, as shown in **Figure 1D**. In Round 1, we collected the top 0.2% of expressing and biotinylated cells, or ~5.9E5 cells. In Rounds 2 and 3, 5.9E4 and 2.0E4 cells were collected, respectively, making up <0.1% of the previously enriched libraries.

Rounds 4 and 5 were negative selections to remove LOV-Turbo mutants with activity in the dark state. Yeast populations were incubated with 50 μM biotin for 4 hours in the dark at 30 °C. Rectangular gates were drawn on the bottom right corner of the AF647 by SA-PE plots. 5.0E5 (2.5%) and 1.8E5 (0.8%) cells were collected, respectively, in Rounds 4 and 5. In Rounds 6 and 7, we performed positive selection at 37 °C rather than 30 °C, to select for mutants with good expression and stability at this higher temperature (the temperature used for applications in mammalian cells). Yeast cells were incubated with 1 μM biotin for 1.5 hours. Rectangular gates were drawn on the top right corner of AF647 by SA-PE plots. Rounds 6 and 7 collected 1.1E4 and 1.5E3 cells, respectively, making up <0.1% of the previously enriched libraries.

### Yeast cell culture

S. cerevisiae strain EBY100 was propagated at 30 °C in SDCAA media supplemented with 20 mg/L tryptophan. Competent yeast cells were transformed with yeast-display plasmid pCTCON2 following the Frozen EZ Yeast Transformation II protocol. Successful transformants were selected in SDCAA plates and propagated in SDCAA media. To induce protein expression, yeasts were inoculated from saturated cultures in 10% SD/GCAA (SDCAA with 90% of dextrose replaced with galactose) or biotin-depleted media (BDM: 1.7 g/L YNB-biotin, 5 g/L ammonium sulfate, 18g/L galactose, 2 g/L dextrose, 5g/L casamino acid, 0.5 nM biotin) overnight from a 1:20-1:100 dilution.

### LOV-Turbo labeling in the yeast cytosol

To collect Western blot samples of BY4741 *S. cerevisiae* in **Fig. 3C**, competent BY4741 were generated and transformed following the Frozen EZ Yeast Transformation II protocol. Yeast cells were plated on leucine-depleted plates (supplemented minimal medium SMM: 6.7 g/L Difco nitrogen base, 20 g/L dextrose, 0.69 g/L CSM-Leu (Sunrise Science Products), with 20 g/L agar) to select for successful transformants at 30 °C. Colonies were then cultured and passaged in SMM. Saturated cultured were then inoculated in 10% D/G SMM (SMM with 90% of dextrose replaced with galactose) to induce expression of the transgene overnight. 800 μl of cultures normalized OD600 to 7 were then pelleted and resuspended in DPBS with or without 100 μM biotin at 30 °C for 6 hours rotating in the dark or exposed to light as indicated. After, samples were kept on ice to stop labeling and washed with DPBS twice. Cells were resuspended with 150 μl of water and 150 μl of 0.6 M NaOH was added. Samples were incubated at room temperature for 5 minutes and then pelleted. 30 μl of 2x Protein loading buffer was used to resuspend cells and boiled for 5 minutes at 95 °C. 15 μl of PBS was then added to dilute the samples.

### LOV-Turbo labeling in *E. coli* cytosol

To collect Western blot samples of BL21 E. coli in **Fig. 3B**, BL21 containing with the indicated constructs were induced with 1mM IPTG overnight in LB media containing selection marker ampicillin. 800 μL of cultures normalized OD600 to 3 were pelleted and resuspended in DPBS with 100 μM biotin where indicated. Samples were incubated for 4 hours at 37 °C, rotating while covered by aluminum foil in the dark, or exposed to light. Bacteria were then pelleted and lysed in 30 μL of 6x protein loading buffer and boiled for 5 minutes at 95 °C. Samples were then diluted with an additional 50 μL of RIPA buffer.

### LOV-Turbo labeling in rat cortical neuron cultures

All procedures were approved and carried out in compliance with the Stanford University Administrative Panel on Laboratory Animal Care, and all experiments were performed in accordance with relevant guidelines and regulations. Prior to dissection, plates were coated with 0.001% (w/v) poly-L-ornithine (Sigma-Aldrich) in DPBS (Gibco) at room temperature overnight, washed twice with DPBS, and subsequently coated with 5 μg/mL of mouse laminin (Gibco) in DPBS at 37°C overnight. Cortical neurons were extracted from embryonic day 18 Sprague Dawley rat embryos by dissociation in HBSS (Gibco) supplemented with 1 mM D-glucose and 10 mM HEPES pH 7.2 (Thermo Life Sciences). Cortical tissue was digested in papain according to the manufacturer’s protocol (Worthington), then plated onto 0.1 mm thick glass coverslips in neuronal plating medium at 37°C under 5% CO2. The neuronal plating medium is 1:1 Advanced DMEM (Gibco): Neurobasal (Gibco), supplemented with 2% (v/v) B27 supplement (Life Technologies), 5% (v/v) fetal bovine serum, 1% GlutaMAX (Gibco), 1% penicillin-streptomycin (VWR, 5 units/mL penicillin, 5 ug/mL streptomycin), 0.1% (v/v) β-mercaptoethanol, 5 ng/mL recombinant human GDNF (Gibco), and 5 μM TRO19622 (Tocris). Every 3 days after plating, half of the media was removed from each well and replaced with neuronal growth medium. The neuronal growth medium is Neurobasal supplemented with 2% (v/v) B27 supplement, 1% GlutaMAX, 50 units/mL penicillin, and 50 μg/mL streptomycin. At 3 days *in vitro* (DIV), 450 μL media was removed from each well and replaced with 500 μL complete neurobasal media supplemented with 10 μM 5-fluorodexoyuridine (FUDR, Sigma-Aldrich) to inhibit glial cell growth. On DIV 6, each well was infected with 2.5e6 vg of concentrated AAV1/2 along with a media change. Neurons were wrapped in aluminum foil and were allowed to express for an additional 5-6 days in the incubator. For **Fig. 4A**, cells were treated with a final concentration of 50 μM biotin and 500 nM MitoTracker (Thermo Fisher, M7512), and exposed to 470 nm blue LED light for 2 hours in the incubator. Cells were fixed and stained for Immunofluorescence detection as described above. For **Fig. 3D and Supplementary Fig. 3C**, cells were treated with a final concentration of 100 μM biotin and exposed to 470 nm blue LED for 30 minutes in the incubator. Cells were lysed or fixed for Western blot analysis or Immunofluorescence detection as described above.

### LOV-Turbo labeling in mouse brain

All animal procedures were performed in accordance with Stanford’s IACUC policies. 8-10 week-old C57BL/6J mice (Jackson Laboratory Strain 000647) were used for all experiments. While under 2-3% isoflurane anesthesia, mice were stereotaxically injected with 500 nL of AAV1/2 in the primary motor cortex (M1) at the following coordinates relative to bregma: anterior +0.62, lateral −1.5, ventral −1.0 mm. Mice were allowed to recover for 14 days. Mice were then placed under anesthesia, injected with 0.5 μL 10 mM biotin at the same stereotaxic coordinates, and implanted with a 200 μm diameter optical fiber (Thorlabs) ~100 μm above the injection site. A subset of mice also received blue light stimulation through the optical fibers using a patch cable and 470 nm fiber-coupled LED (Thorlabs; 5 mW/cm^2^ of 470nm light measured at the fiber tip, then 10 Hz pulses, 10 ms pulse width, 10% duty cycle) for 1.5 hours. After light treatment, both experimental and control groups of mice were sacrificed for tissue collection.

Mice were deeply anesthetized using 1 mL intraperitoneally delivered 2.5% Avertin. They were then quickly perfused with 10 mL ice-cold PBS, and their brains were immediately dissected and cut in a ~1 mm coronal section using a brain block and razor blades on ice. The M1 cortex was micro-dissected from the coronal slice and placed in 200 μL of ice-cold buffer, 50 mM Tris/HCl, pH 7.5, 150 mM NaCl, 1 mM EDTA supplemented with protease inhibitor cocktail and PMSF. Each sample contained M1 dissected from five ~1 mm coronal sections. A tissue homogenizer was then used to process the tissue. The samples were lysed by adding 1 mL of ice-cold buffer containing 0.2% SDS, 1% TrixonX-100, and 1% sodium deoxycholate. Samples were incubated with benzonase (50 U/mL) at 4 °C for 30 minutes and briefly sonicated for 30 seconds. Lysates were cleared via centrifugation at 20,000 rcf at 4 °C for 30 minutes, and then processed for Western blot analysis as described above.

### AAV generation

Mito-LOV-Turbo, mito-TurboID, LOV-Turbo-NES, and TurboID-NES were cloned into an AAV1/2 vector backbone using Gibson overlap assembly. HEK 293T cells (3 T150 flasks maintained in DMEM + 10% FBS) were transfected with 15.6 μg of plasmid, 13.1 μg of AAV1 serotype plasmid, 13.1 μg of AAV2 serotype plasmid, 31.2 μg of DF6 helper plasmid, and 390 μL of PEI. 48 hours later, the media was aspirated, and the cells were collected using a cell scraper in DPBS. The cells were pelleted, the DPBS removed, and then the pellet was resuspended in 20 mL of 100 mM NaCl, 20 mM Tris (pH 8.0). A final solution of 0.5% sodium deoxycholate and 50 U/mL of benzonase nuclease (Sigma) was added to the solution, and the mixture was incubated at 37 °C for 1 hour. Cellular debris was removed by an additional centrifuge step, and the supernatant was then run through a heparin column (Cytiva). The column was washed with increasing salt concentrations (10 mL 100 mM, 1 mL 200 mM, and 1 mL 300 mM NaCl in 20 mM Tris, pH 8.0). The virus was then eluted using 1.5 mL 400 mM, 3 mL 450 mM, and 1.5 mL 500 mM NaCl in 20 mM Tris, pH 8.0. The virus was concentrated using an Amicon Ultra 15 mL centrifugal filter unit with a 100,000 molecular weight cutoff a final volume of 100 μL was reached. The viruses were titrated using SybrGreen qPCR mastermix using primers detecting the WPRE (AAV1/2-Syn-Mito-LOV-Turbo: 3.35E9 vg/μL; AAV1/2-Syn-Mito-TurboID: 4.03E9 vg/μL; AAV1/2-CAG-LOV-Turbo-NES: 1.26E10 vg/μL; AAV1/2-CAG-TurboID-NES: 2.55E10 vg/μL)

### BRET activation of LOV-Turbo

To generate samples used in **Fig. 4G,** cells were transiently transfected with V5-NanoLuc-LOV-Turbo-NES fusion as described and induced to express overnight with doxycycline in the dark. Cells were then treated to 100 μM biotin and either kept in the dark, stimulated with 10 minutes of light, or stimulated for 10 minutes or 1 hour by BRET through activation of NanoLuc’s bioluminescence with its substrate furimazine (1:100 dilution of Promega nano-Glo Live Assay). After treatment, cells were washed in cold DPBS twice and lysed in RIPA buffer containing protease inhibitor cocktail. Samples were then combined with protein loading buffer and boiled at 95 °C for 5 minutes for gel loading.

To generate samples used in **Fig. 4I**, cells were transiently co-transfected with CCR6-NanoLuc-V5 and Myc-Arrestin-LOV-Turbo and induced to express overnight with doxycycline in the dark. Cells were then treated to 100 μM biotin and either kept in the dark, stimulated with 10 minutes of light, or stimulated with 1 hour of furimazine and with or without CCR6 activation by addition of its substrate CCL20 (0.2 μg/mL). After treatment, cells were washed in cold DPBS twice and lysed in RIPA buffer containing protease inhibitor cocktail. Samples were then combined with protein loading buffer and incubated at 37 °C for 10 minutes for gel loading.

### Analysis of LOV-Turbo reversibility

To generate samples in **Fig. 3A**, HEK 293T cells expressing LOV-Turbo-NES were induced to express overnight with doxycycline. Cells were either treated to neither light nor biotin (A), 1 hour of 100 μM biotin (B), 1 hour of light stimulation (C), 1 hour of biotin and light stimulation (D), 1 hour of biotin with 15 minutes of light in the first 15 minutes (E), or 15 minutes of biotin and light stimulation (F). Following treatments, cells were washed twice with cold DPBS and lysed in RIPA buffer containing protease inhibitor cocktail and processed for Western blot analysis.

### Analysis of LOV-Turbo light specificity

To generate samples in **Supplementary Fig. 2E,** HEK 293T cells expressing LOV-Turbo-NES were induced to express overnight with doxycycline. Cells were then incubated in 100 μM biotin media and either treated to no light, blue-light stimulation (AMUZA), or red-light stimulation (HQRP, #884667920839, 630nm, 14W) for 30 minutes. Cells were then washed twice with cold DPBS, lysed in RIPA buffer containing protease inhibitor cocktail and processed for Western blot.

### Analysis of LOV-Turbo turn-on and turn-off kinetics

To generate samples in **Supplementary Fig. 3A** and test the turn-on kinetics of LOV-Turbo, HEK 293T cells expressing LOV-Turbo-NES were induced to express overnight with doxycycline. Cells were treated to 5 minutes of blue light with either 0-, 1-, 2-, or 3-minutes delay in 100 μM biotin addition. A non-expressing cell was used as a control and treated with 5 minutes of light and biotin. Cells were then washed twice with cold DPBS, lysed in RIPA buffer containing protease inhibitor cocktail and processed for Western blot. Labeling intensity was quantified by the intensity of the streptavidin lane signal and normalized by the V5 lane signal.

To generate samples in **Supplementary Fig. 3B** and test the turn-off kinetics of LOV-Turbo, HEK 293T cells expressing LOV-Turbo-NES were induced to express overnight with doxycycline. Cells were treated with the following conditions: 15 minutes of blue light, 15 minutes of 100 μM biotin, 15 minutes of blue light followed by 1-minute interval then 15 minutes of biotin, 15 minutes of blue light followed immediately by 15 minutes of biotin, or 15 minutes of blue light and biotin. Non-expressing cells were used as a control and treated with 15 minutes of blue light and biotin. Cells were then washed twice in cold DPBS, lysed in RIPA buffer containing protease inhibitor cocktail and processed for Western blot.

### Analysis of biotin washout

To generate samples in **Supplementary Fig. 3E** and test the efficiency of biotin washout, HEK 293T cells expressing LOV-Turbo-NES were induced to express overnight with doxycycline. Cells were preincubated with 100 μM biotin for 10 minutes then either collected immediately for Western blotting or washed once, twice, or 5 times before replacing with media followed by light stimulation for 30 minutes. The remaining samples were then lysed and processed for Western blotting.

### Analysis of spatial specificity by light activation

To generate samples used in **Fig 4B**, stable LOV-Turbo-NES expressing cells were plated on cell culture chamber slides (Southern Labware 230118) and induced with doxycycline overnight. Half of the wells were covered with black tape on the top and bottom to obstruct light. Cells were then labeled with 5 minutes of light in 100 μM biotin media. Cells were fixed, permeabilized, and stained with DAPI, anti-V5 antibody, and neutravidin.

### LOV-Turbo labeling for pulse-chase proteomics

To generate samples for **Fig. 4C and Supplementary Fig. 3F-G**, HEK 293T cells expressing LOV-Turbo-NLS plated on human fibronectin-covered glass coverslips were induced to express overnight with doxycycline. Cells were labeled with 100 μM biotin and blue light for 10 minutes followed by 0, 2, 4 or 8 hours chase in the dark in fresh media. Cells were then fixed and processed for immunofluorescence to detect LOV-Turbo-NLS labeling and localization by staining with primary antibody anti-V5 followed by staining with anti-mouse-AlexaFluor488, DAPI, and neutravidin-AlexaFluor647.

To generate samples for **Fig. 4E**, HEK 293T cells expressing LOV-Turbo-NES plated 10 cm cell culture dishes were induced to express overnight with doxycycline. Cells were labeled with 100 μM biotin and blue light for 30 minutes followed by 0- or 2-hour chase in the dark in fresh media at either 37 °C or 42 °C. Cells were collected in 2 ml DPBS. 50 μl was removed and lysed in RIPA buffer containing protease inhibitor cocktail to generate the whole cell lysate samples. The remainder was processed for nuclear fractionation. The nuclear fraction was then lysed in 500 μl RIPA buffer containing protease inhibitor cocktail and benzonase. 50 μl were removed to generate the nuclear lysate samples. The remainder was processed for streptavidin enrichment. All samples from the whole cell lysate, nuclear lysate, and streptavidin enrichment were analyzed by Western blotting using streptavidin, anti-importin-α antibody, and anti-HSP70 antibody.

To generate samples for **Supplementary Fig. 4A**, HEK 293T cells stably expressing LOV-Turbo-ERM or LOV-Turbo-NES plated CELLSTAR® OneWell Plate (Greiner Bio-One, 670180) were induced to express overnight with doxycycline. Cells were labeled with 100 μM biotin and blue light for 30 minutes followed by a 2-hour chase in the dark in fresh media with or without 2 μg/ml tunicamycin. Cells were collected in 30 mL DPBS and washed three times and resuspended in 750 μL. 50 μL was removed and lysed in RIPA buffer containing protease inhibitor cocktail to generate the whole cell lysate samples. The remainder of the whole cell lysate was processed for streptavidin enrichment. The remainder of the cells in DPBS was processed for nuclear fractionation. The cytosolic fraction was also collected in the process of nuclear fractionation. The nuclear fraction was then lysed in 500 μL RIPA buffer containing protease inhibitor cocktail and benzonase. 20 μL were removed to generate the nuclear lysate samples. The remainder of the nuclear fraction was processed for streptavidin enrichment. All samples from the whole cell lysate, cytosolic lysate, nuclear lysate, and streptavidin enrichment of the nuclear and whole cell lysate were analyzed by Western blotting using streptavidin, anti-p84 antibody, anti-LaminB1 antibody, anti-calnexin antibody, anti-ATF6 antibody, anti-V5 antibody, and anti-GAPDH antibody.

To generate samples for **Supplementary Fig. 5A**, HEK 293T cells expressing LOV-Turbo-ERM plated 10 cm cell culture dishes were induced to express overnight with doxycycline. Cells were labeled with 100 μM biotin and blue light for 1 hour followed by an 8-hour chase in the dark in fresh media with or without 1 μM FCCP in duplicates. Cells were collected in 2 ml DPBS. 50 μL was removed and lysed in RIPA buffer containing protease inhibitor cocktail to generate the whole cell lysate samples. The remainder was processed for mitochondrial fractionation. The mitochondrial fraction was then lysed in 500 μL RIPA buffer containing protease inhibitor cocktail. 50 μL were removed to generate the mitochondria lysate samples. The remainder was processed for streptavidin enrichment. All samples from the whole cell lysate, mitochondria lysate, and streptavidin enrichment were analyzed by Western blotting using streptavidin, anti-V5 and anti-SDHA antibody.

### Streptavidin enrichment of LOV-Turbo labeled material

Streptavidin enrichment was performed following the protocol in Cho et al.^7^ Streptavidin beads were washed with 1 ml RIPA buffer twice before adding to the cell lysate and incubated overnight at 4 °C overnight or 1 hour at room temperature, rotating. Beads were then washed thrice with 1 ml RIPA buffer, followed by washes of 1 M KCl, 0.1 M Na2CO2, and 2 M urea in 10 mM Tris-HCl (pH 8). Beads were then washed with 1 ml RIPA three more times. Then, proteins were eluted from beads using 3x protein loading buffer containing 2 mM biotin and 20 mM DTT at 95 °C for 5 minutes.

### Nuclear fractionation

Nuclear fractionations were performed following the protocols and suggestions published by Gagnon, KT et al.^54^, and Senichkin, VV. et al.^55^ Cells were collected in DPBS with some aliquoted for whole cell lysate samples then pelleted at 500 rcf at 4 °C for 2 minutes. 1 mL of hypotonic lysis buffer (HLB, 20mM Tris (pH7.5), 5mM KCl, 3mM MgCl2, 10% glycerol, 0.5% NP-40, and protease inhibitor cocktail) was used to resuspend 75 mg of cells and incubated on ice for 10 minutes. Cells were briefly vortexed and centrifuged at 500 rcf for 5 minutes at 4°C. 870 μL of the supernatant was transferred to a new tube and combined with 25 μL of 5 M NaCl to generate the cytoplasmic fraction. The remaining supernatant was discarded, and the pellet was resuspended in 1 mL HLB. The nuclear fraction was pelleted at 500 rcf for 2 minutes and then washed in cold isotonic wash buffer (IWB, 20 mM Tris-HCl (pH 7.5), 100 mM KCl, 3 mM MgCl2, 10% glycerol, 0.5 mM DTT, 0.5% NP-40 and protease inhibitor cocktail). This washing step with the IWB was performed once more. Then, the nuclear pellet was lysed in RIPA buffer containing benzonase at 1000 U/mL (Millipore).

### Mitochondrial fractionation

Mitochondria fractionations were performed using the Pierce Mitochondria Isolation Kit for Tissue with the following modifications. Cells were collected in DPBS. Prior to fractionation, a subset of cells was aliquoted to generate the whole cell lysate samples. Cells were pelleted at 700 rcf for 2 minutes at 4 °C. The supernatant was removed, and the cell pellet was resuspended in 800 μL of Reagent A supplemented with a protease inhibitor cocktail for 2 minutes. The cells were then lysed using a Dounce tissue grinder (Kimble) 15 times and then transferred into a 2 mL tube. 800 μL of Reagent C supplemented with protease inhibitor cocktail was added. The grinder was washed with 200 μL of Reagent A which was then added back to the cells. The cells were then centrifuged at 700 rcf for 10 minutes at 4 °C and the supernatant was transferred to a new tube. This step was repeated until a pellet was no longer visible following centrifugation. Proteinase K was added at 10 U/mL for 20 minutes at 4 °C to remove proteins on the outside of the mitochondria, then PMSF was supplemented at 1 mM to inhibit it. The mitochondrial fraction was pelleted at 3000 rcf for 15 minutes and the pellet was washed twice with Reagent C, centrifuging after each wash at 12000 rcf for 5 minutes. Finally, the mitochondrial pellet was lysed in RIPA lysis buffer.

### Sample preparation for mass spectrometry

To generate samples for ERM to mitochondria translocation profiling, cells stably expressing LOV-Turbo-ERM or LOV-Turbo-NES were plated on 127.5 x 85.5 mm plates and induced with doxycycline overnight. Cells were treated with 100 μM biotin and stimulated with light or kept in the dark for 1 hour. Biotin media was washed out with DPBS three times and then replaced with fresh media containing 1 μM FCCP, 100 nM thapsigargin, or no drug for 8 hours. Cells were then collected and fractionated for their mitochondria and lysed in 500 μl RIPA buffer containing protease inhibitor cocktail. Mitochondrial lysates were flash-frozen with liquid nitrogen and kept at −80 °C until ready to proceed to streptavidin enrichment. For each channel, samples collected from 2 plates were combined. Combined 1 ml of mitochondrial lysate were then incubated with 50 μl of streptavidin beads for enrichment and incubated with the beads overnight at 4 °C. After washing, 2.5% of each sample was removed to be eluted with protein loading buffer and checked by silver stain (Pierce) for the presence of sufficient proteins. The remaining beads were washed twice with 75 mM NaCl in 50 mM Tris-HCl (pH 8) and then flash-frozen in 50 μl of the buffer.

To generate samples for ERM to nucleus translocation profiling, cells stably expressing LOV-Turbo-ERM or LOV-Turbo-NES were plated on 127.5 x 85.5 mm plates and induced with doxycycline overnight. Cells were treated with 100 μM biotin and stimulated with light or kept in the dark for 30 minutes. Biotin media was replaced with fresh media containing 2 μg/ml tunicamycin for 2 hours. Cells were then collected and fractionated for their nucleus and lysed in 500 μL RIPA buffer containing protease inhibitor cocktail. For each channel samples collected from 2 plates were combined. Clarified lysates were incubated with 35 μL of streptavidin beads for enrichment and incubated for 1 hour at room temperature. After washing, 2.5% of each sample was removed to be eluted with protein loading buffer and checked by silver stain (Pierce) for the presence of sufficient proteins. The remaining beads were washed twice with 75 mM NaCl in 50 mM Tris-HCl (pH 8) and then flash-frozen in 50 μl of the buffer.

### On-bead trypsin digestion of biotinylated proteins

Samples collected and enriched with streptavidin magnetic beads were washed twice with 200 μL of 50mM Tris-HCl buffer (pH 7.5), transferred into new 1.5 mL Eppendorf tubes, and washed 2 more times with 200 μL of 50mM Tris (pH 7.5) buffer. Samples were incubated in 0.4 μg trypsin in 80 μL of 2M urea/50mM Tris buffer with 1 mM DTT, for 1 h at room temperature while shaking at 1000 rpm. Following pre-digestion, 80 μL of each supernatant was transferred into new tubes. Beads were then incubated in 80 uL of the same digestion buffer for 30 min while shaking at 1000rpm. Supernatant was transferred to the tube containing the previous elution. Beads were washed twice with 60 μL of 2M urea/50mM Tris buffer, and these washes were combined with the supernatant. The eluates were spun down at 5000 × g for 30 seconds and the supernatant was transferred to a new tube. Samples were reduced with 4 mM DTT for 30 min at room temperature, with shaking. Following reduction, samples were alkylated with 10mM iodoacetamide for 45 min in the dark at room temperature. An additional 0.5 μg of trypsin was added and samples were digested overnight at room temperature while shaking at 700 × g. Following overnight digestion, samples were acidified (pH < 3) with neat formic acid (FA), to a final concentration of 1% FA. Samples were spun down and desalted on C18 StageTips as previously described. Eluted peptides were dried to completion and stored at −80 °C.

### TMT labeling and fractionation

Desalted peptides were labeled with TMTpro (18-plex) reagents (ThermoFisher Scientific). Peptides were resuspended in 80 μL of 50 mM HEPES and labeled with 20 μL 25 mg/mL TMTpro18 reagents in ACN. Samples were incubated at RT for 1 h with shaking at 1000 rpm. TMT reaction was quenched with 4 μL of 5% hydroxylamine at room temperature for 15 minutes with shaking. TMT-labeled samples were combined, dried to completion, reconstituted in 100 μL of 0.1% FA, and desalted on StageTips.

TMT labeled peptide sample was fractionated by basic reverse phase (bRP) fractionation. StageTips packed with two disks of SDB-RPS (Empore) material. StageTips were conditioned with 100 μL of 100% MeOH, followed by 100 μL 50% MeCN/0.1% FA and two washes with 100 μL 0.1% FA. Peptide samples were resuspended in 200 μL 1% FA (pH<3) and loaded onto StageTips. 8 step-wise elutions were carried out in 100 μL 20 mM ammonium formate buffer with increasing concentration of 5%, 7.5%, 10%, 12.5%, 15%, 20%, 25%, and 45% MeCN. Eluted fractions were dried to completion.

### Liquid chromatography and mass spectrometry

All peptide samples were separated with an online nanoflow Proxeon EASY-nLC 1200 UHPLC system (Thermo Fisher Scientific) and analyzed on an Orbitrap Exploris 480 mass spectrometer (Thermo Fisher Scientific). In this set up, the LC system, column, and platinum wire used to deliver electrospray source voltage were connected via a stainless-steel cross (360 mm, IDEX Health & Science, UH-906x). The column was heated to 50 °C using a column heater sleeve (Phoenix-ST). Each sample was injected onto an inhouse packed 27 cm x 75 μm internal diameter C18 silica picofrit capillary column (1.9 mm ReproSil-Pur C18-AQ beads, Dr. Maisch GmbH, r119.aq; PicoFrit 10 μm tip opening, New Objective, PF360-75-10-N-5). Mobile phase flow rate was 200 nL/min, comprised of 3% acetonitrile/0.1% formic acid (Solvent A) and 90% acetonitrile/0.1% formic acid (Solvent B). The 154-min LC–MS/MS method used the following gradient profile: (min:%B) 0:2;2:6; 122:35; 130:60; 133:90; 143:90; 144:50; 154:50 (the last two steps at 500 nL/min flow rate). Data acquisition was done in the data-dependent mode acquiring HCD MS/MS scans (r = 45,000) after each MS1 scan (r = 60,000) on the top 12 most abundant ions using a normalized MS1 AGC target of 100% and an MS2 AGC target of 50%. The maximum ion time utilized for MS/MS scans was 120 ms; the HCD-normalized collision energy was set to 32; the dynamic exclusion time was set to 20 s, and the peptide match and isotope exclusion functions were enabled. Charge exclusion was enabled for charge states that were unassigned, 1 and >6.

### Analysis of mass spectrometry data (peptide level, protein level)

Mass spectrometry data was processed using Spectrum Mill v 7.11 (proteomics.broadinstitute.org). For all samples, extraction of raw files retained spectra within a precursor mass range of 600-6000 Da and a minimum MS1 signal-to-noise ratio of 25. MS1 spectra within a retention time range of +/- 45 s, or within a precursor m/z tolerance of +/- 1.4 m/z were merged. MS/MS searching was performed against a human Uniprot database with a release date of December 28, 2017. Digestion parameters were set to “trypsin allow P” with an allowance of 4 missed cleavages. The MS/MS search included fixed modification of carbamidomethylation on cysteine. TMTpro18 was searched using the full-mix function. Variable modifications were acetylation and oxidation of methionine. Restrictions for matching included a minimum matched peak intensity of 30% and a precursor and product mass tolerance of +/- 20 ppm.

Peptide spectrum matches (PSMs) were validated using a maximum false discovery rate (FDR) threshold of 1.2% for precursor charges 2 through 6 within each LC-MS/MS run. Protein polishing autovalidation was further applied to filter the PSMs using a target protein score threshold of 9. TMTpro18 reporter ion intensities were corrected for isotopic impurities in the Spectrum Mill protein/peptide summary module using the afRICA correction method which implements determinant calculations according to Cramer’s Rule. We required fully quantified unique human peptides for protein quantification. We used the Proteomics Toolset for Integrative Data Analysis (Protigy, v1.0.4, Broad Institute, https://github.com/broadinstitute/protigy) to calculate moderated *t*-test *P* values for regulated proteins.

### Data Accessibility

The original mass spectra, spectral library, and the protein sequence database used for searches have been deposited in the public proteomics repository MassIVE (http://massive.ucsd.edu) and are accessible at ftp://MSV000090683@massive.ucsd.edu when providing the dataset password: allostery. If requested, also provide the username: MSV000090683. These datasets will be made public upon acceptance of the manuscript.

### Analysis of mass spectrometry data

To check the specificity of ERM labeling and nuclear fractionation, nuclear fraction sample from LOV-Turbo ERM labeling under basal condition was analyzed compared to (1) no-light, (2) omit-fractionation (3) LOV-Turbo-NES labeling. We ranked the MS-detected proteins by fold-change (FC) values and performed GOCC term enrichment analysis by g:Profiler^56^ (**Supplementary Fig. 4E-G**). For (1), cutoffs were set by ROC analysis using a list of nuclear proteins as true positives and mitochondrial proteins as false positives (**Supplementary Fig. 4E**). This resulted in a Log_2_FC cutoff at 3.014, and we also applied a p-value cutoff of 0.05, to generate a list of 2761 proteins. **Supplementary Fig. 4E** shows nuclear terms enriched in GOCC analysis. For (2), cutoffs were set by ROC analysis using a list of a list of nuclear proteins as true positives and mitochondrial proteins as false positives (**Supplementary Fig. 4F**). This resulted in a Log_2_FC cutoff at −0.861, and we also applied a p-value cutoff of 0.05, to generate a list of 1799 proteins. **Supplementary Fig. 4F** shows nuclear terms enriched in GOCC analysis. For (3), cutoffs were set by ROC analysis using a list of ER membrane proteins as true positives and mitochondrial proteins as false positives (**Supplementary Fig. 4G**). This resulted in a Log_2_FC cutoff at 0.951, and we also applied a p-value cutoff of 0.05, to generate a list of 1019 proteins. **Supplementary Fig. 4G** shows ER terms enriched in GOCC analysis.

To filter ER membrane-associated proteins that traffick to the nucleus under tunicamycin-induced ER stress conditions (**Supplementary Fig. 4H** that shows the flow chart for data analysis), we ranked the MS-detected proteins from LOV-Turbo-ERM labeling under stress conditions by four different fold-change values: (1) LOV-Turbo-ERM labeling under basal condition, (2) no-light condition, (3) LOV-Turbo-NES labeling under stress condition, and (4) omit nuclear fractionation (whole cell lysates) under stress condition. For (1), cutoffs were set at a Log_2_FC value of 0.1, a p-value cutoff of 0.05, and unique peptides ≥ 2. This generated a list of 131 proteins. For (2)-(4), cutoffs were set at a Log_2_FC value of 2.3, 0.82, and - 2.7 respectively and a p-value cutoff of 0.05. The filtered 29 proteins are listed in **Supplementary table 2**.

To identify ER membrane-associated proteins that traffick to the mitochondrion under basal conditions (**Supplementary Fig. 5E** that shows the flow chart for data analysis), we ranked the MS-detected proteins by three different fold-change values: (1) LOV-Turbo-ERM labeling compared to FCCP treatment during chase, (2) LOV-Turbo-ERM labeling compared to no-light condition, and (3) LOV-Turbo-ERM labeling compared to LOV-Turbo-NES labeling. For (1), cutoffs were set at a Log_2_FC value of 0 and a p-value cutoff of 0.05. This generated a list of 595 proteins. For (2), cutoffs were set at a 10% false discovery rate and p-value < 0.05, using a list of nuclear proteins as false positives. This resulted in a list of 539 proteins. For (3), cutoffs were set by ROC analysis using a list of ER membrane proteins as true positives and nuclear proteins as the false positives (**Supplementary Fig. 5D**). This resulted in a Log_2_FC cutoff at −0.746, and we also applied a p-value cutoff of 0.05, to generate a list of 747 proteins. The intersection of the three lists resulted in 16 proteins (**Supplementary Table 3**).

To identify ER membrane-associated proteins that traffick to the mitochondrion under thapsigargin-induced ER stress conditions (**Supplementary Fig. 5E** that shows the flow chart for data analysis), we ranked the MS-detected proteins by three different fold-change values: (1) LOV-Turbo-ERM labeling compared to thapsigargin treatment during chase, (2) LOV-Turbo-ERM labeling compared to no-light condition, and (3) LOV-Turbo-ERM labeling compared to LOV-Turbo-NES labeling. For (1), cutoffs were set at a Log_2_FC value of 0 and a p-value cutoff of 0.05. This generated a list of 963 proteins. For (2), cutoffs were set at a 10% false discovery rate and p-value < 0.05, using a list of nuclear proteins as the false positives. This resulted in a list of 692 proteins. For (3), cutoffs were set by ROC analysis using a list of ER membrane proteins as true positives and nuclear proteins as the false positives (**Supplementary Fig. 5D**). The resulted in a Log_2_FC cutoff at −0.727, and we also applied a p-value cutoff of 0.05, to generate a list of 674 proteins. The intersection of the three lists resulting in 64 proteins (**Supplementary Table 3**).

### Cloning

For cloning, fragments were PCR amplified using Q5 polymerase (NEB) and vectors were digested using restriction enzymes. All fragments were gel purified and ligated using Gibson assembly (NEB), then transformed into stable competent *E. coli* (NEB). LOV-Turbo gene was codon optimized for human via GenSmart™. Below is the complete amino acid sequence of our final optimized LOV-Turbo:

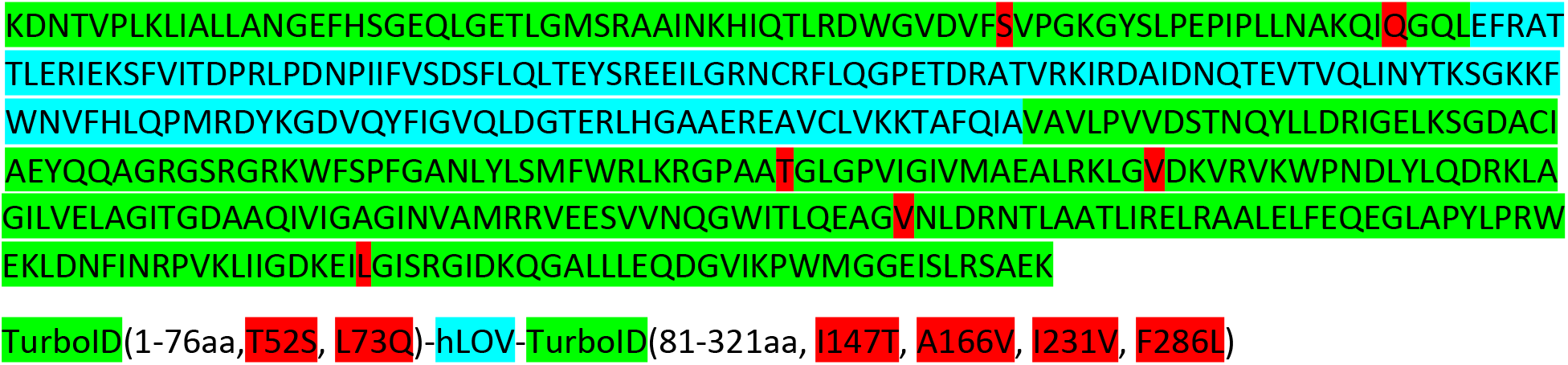

To target LOV-Turbo to various subcellular compartments, we created genetic fusions to a nuclear export signal (NES, C-terminal), nuclear localization signal (NLS, C-terminal), SEC61B protein (to target to ER membrane facing cytosol, C-terminal), COX4I1 mitochondrial targeting sequence (MTS) (to target to the mitochondrial matrix (1) in **Fig. 2D**, N-terminal), COX4I1 MTS-NES (to target to the mitochondrial matrix (2) in **Fig. 2D**, N-terminal for COX4I1 MTS and C-terminal for NES), 3x COX8A MTS-NES (to target to the mitochondrial matrix (3) in **Fig. 2D**, N-terminal for 3xCOX8A MTS and C-terminal for NES), or AKAP1 transmembrane domain (to target to the outer mitochondrial membrane, N-terminal). For mitochondrial matrix targeting, COX4I1 worked the best. Specific amino acid sequences of targeting peptides are given in Supplementary Information under “Genetic constructs used in this study”.

