## Supplementary information for "Engineered allostery in light-regulated LOV-Turbo enables precise spatiotemporal control of proximity labeling in living cells"

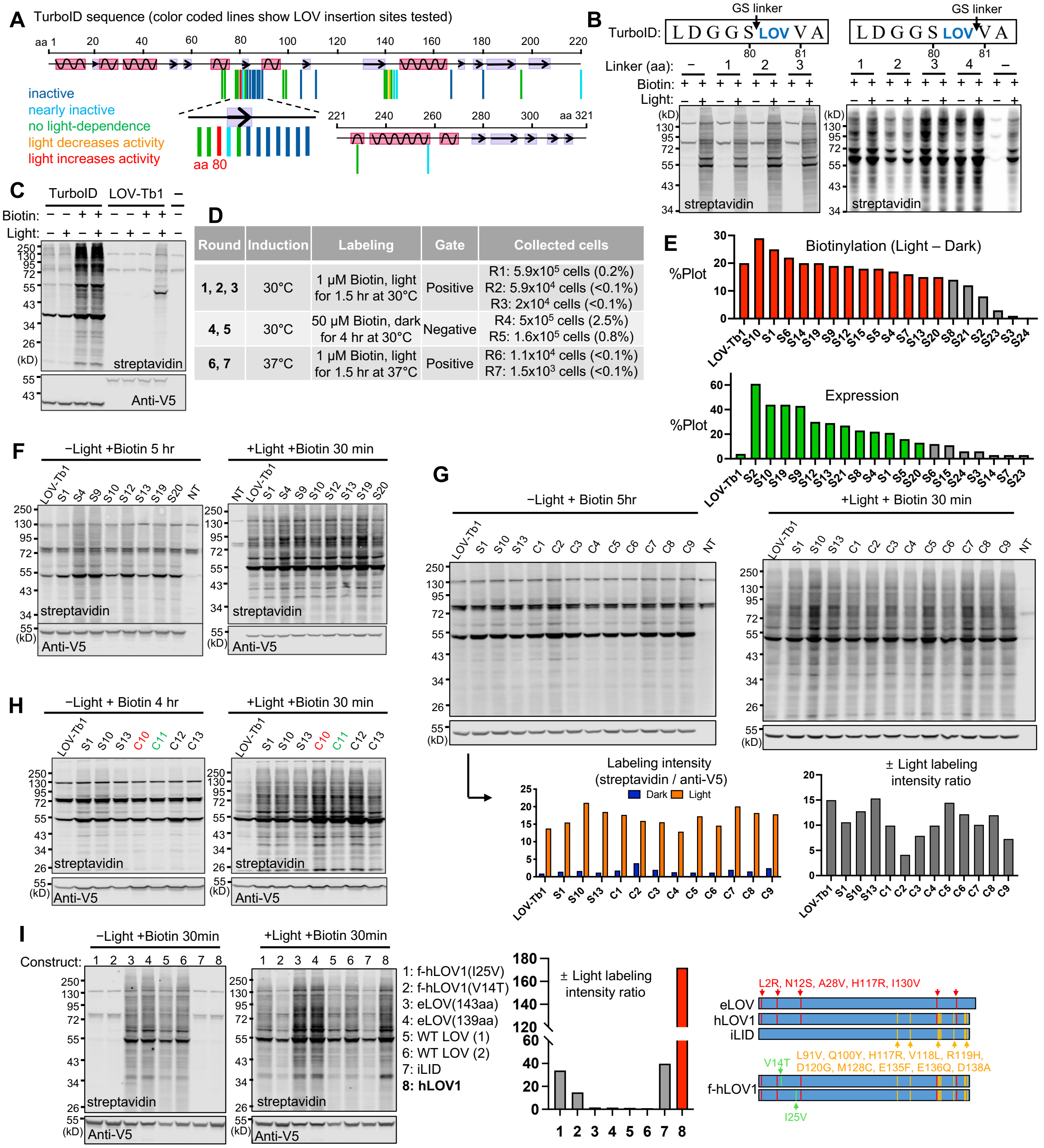


| Round | Induction | Labeling | Gate | Collected cells |
|---|---|---|---|---|
| 1, 2, 3 | 30°C | 1 μM Biotin, light for 1.5 hr at 30°C | Positive | R1: 5.9x$10^5$ cells (0.2%)<br>R2: 5.9x$10^4$ cells (<0.1%)<br>R3: 2x$10^4$ cells (<0.1%) |
| 4, 5 | 30°C | 50 μM Biotin, dark for 4 hr at 30°C | Negative | R4: 5x$10^5$ cells (2.5%)<br>R5: 1.6x$10^5$ cells (0.8%) |
| 6, 7 | 37°C | 1 μM Biotin, light for 1.5 hr at 37°C | Positive | R6: 1.1x$10^4$ cells (<0.1%)<br>R7: 1.5x$10^3$ cells (<0.1%) |

**Supplementary Fig. 1. Additional data related to the design and directed evolution of LOV-Turbo.** (**A**) Domain structure of TurboID showing LOV insertion sites tested and the result, by color. α-helices are pink and β-sheets are purple. (**B**) Testing GS linkers around the 80/81 LOV insertion site. Linkers of various lengths (G, GS, GSG, or GSGS) were tested on either side of the inserted hLOV1. Constructs were expressed in the HEK cytosol with labeling was performed for 30 minutes in light or dark. (**C**) LOV-Turbo1 expression level is lower than that of TurboID. (**D**) A more detailed version of the table in **Fig. 1E**. (**E**) Testing of 19 evolved mutants on the yeast cell surface (mutations listed in **Supplementary Table 1**). Biotinylation activity and expression level of each clone quantified by FACS as described in **Methods**. Labeling time, 1.5 hours. Top-performing clones (colored) were tested in HEK 293T cells in **Fig. 1G**. (**F**) Streptavidin and anti-

V5 blots used to generate data in **Fig. 1G**. (**G**) Screening of 9 combination mutants C1-C9 after labeling for 30 minutes in light or 5 hours in dark. Since L73M increases background while L73Q decreases background in the dark (C9 vs C6, mutations listed in **Supplementary Table 1**), L73Q was selected for further combination testing. (**H**) Streptavidin and anti-V5 blots used to generate data in **Fig. 1H**. (**I**) Testing alternative LOV domains in LOV-Turbo. Constructs contained the indicated LOV domains (f-hLOV1[1, 2], eLOV[3], iLID[4], wild-type AsLOV2, or hLOV1[5]). Mutations relative to wild-type AsLOV2 are shown at right. Full sequences are provided under "Genetic constructs used in this study". Constructs were expressed in the HEK cytosol and labeling was performed for 30 minutes in the presence or absence of light. ±Labeling intensity ratios were quantified after streptavidin blotting of whole cell lysates. Construct 8 is our final optimized LOV-Turbo.

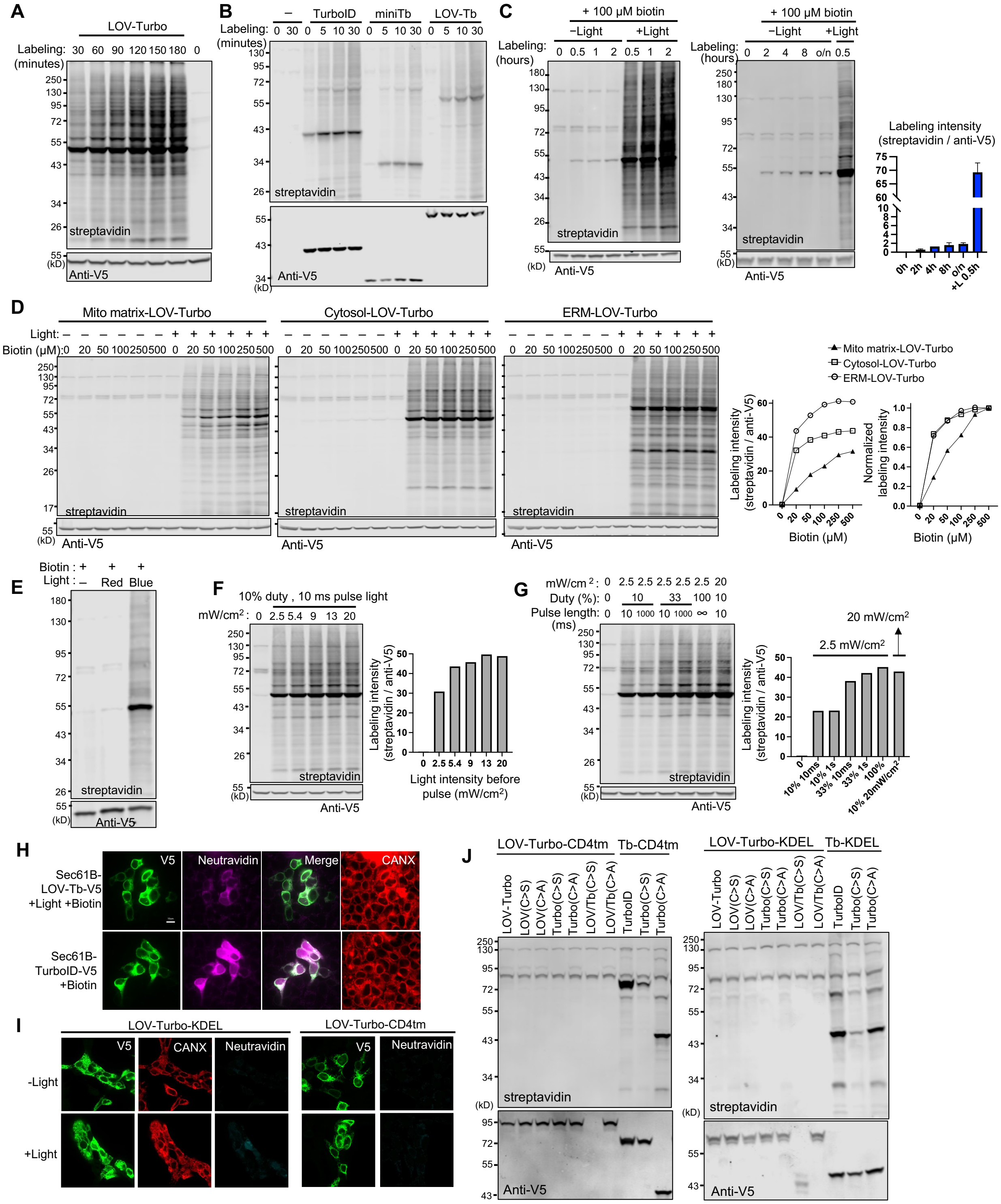


**Supplementary Fig. 2. Additional data related to the characterization of LOV-Turbo.** (**A**) LOV-Turbo labeling time course. Labeling was performed in the HEK cytosol for 0-180 minutes. (**B**) Labeling time course for LOV-Turbo (LOV-Tb), TurboID, and miniTurbo (miniTb) in the HEK cytosol. Labeling was performed for 5-30 minutes. (**C**) LOV-Turbo displays minimal background labeling in the dark. HEK293T cells expressing cytosolic LOV-Turbo were labeled with biotin in the dark for up to 2 hours (left) or up to overnight (right). Lysates were analyzed by blotting with streptavidin and anti-V5 antibody. Relative labeling intensities are quantified for right blot. (**D**) Biotin concentration titration. HEK 293T cells expressing LOV-Turbo in the mitochondrial matrix, cytosol, or ER membrane were exposed to 0-0.5 mM biotin for 30 minutes in light or dark. Mito-LOV-Turbo requires higher biotin concentration than cytosolic LOV-Turbo. Relative labeling intensities quantified from streptavidin and anti-V5 blots. (**E**) LOV-Turbo is activated by blue light but not red light. Cytosolic LOV-Turbo expressed in HEK 293T cells was treated with biotin and light for 30 minutes. (**F**) Testing of various pulsed light sequences for LOV-Turbo activation. HEK 293T cells expressing cytosolic LOV-Turbo were exposed to 10 ms pulses of 0-20 $mW/cm^2$ light every 100 ms (10% duty cycle) for 15 minutes in the presence of biotin. Lysates were analyzed by blotting with streptavidin and anti-V5 antibody. Relative labeling intensity was quantified by streptavidin and anti-V5 blots. (**G**) Same as (F) but cells were exposed to different light pulse lengths (10 ms or 1000 ms) and duty cycles (10%, 33% or 100%). (**H**) LOV-Turbo fused to SEC61B has low activity in mammalian cells. Both LOV-Turbo and TurboID were targeted to ER membrane, facing the ER lumen, via fusion to the ER transmembrane protein SEC61B, and labeling was performed in HEK 293T cells for 30 minutes. Samples were fixed and stained with anti-V5 to detect ligase expression, neutravidin to detect biotinylated proteins, and anti-calnexin antibody to detect the ER. Scale bar, 10µm. (**I**) LOV-Turbo-KDEL and cell surface LOV-Turbo (LOV-Turbo-CD4 transmembrane) are both nearly inactive in mammalian cells. Constructs were expressed in HEK 293T cells and labeling was performed for 1 hour with biotin and light. Samples were fixed and stained as in (H). (**J**) Mutation of cysteines in LOV-Turbo does not restore activity in the ER lumen or on the cell surface. Cys128 in hLOV1 or Cys103 in TurboID were mutated to serine or alanine. Labeling was performed in HEK 293T cells for 1 hour.

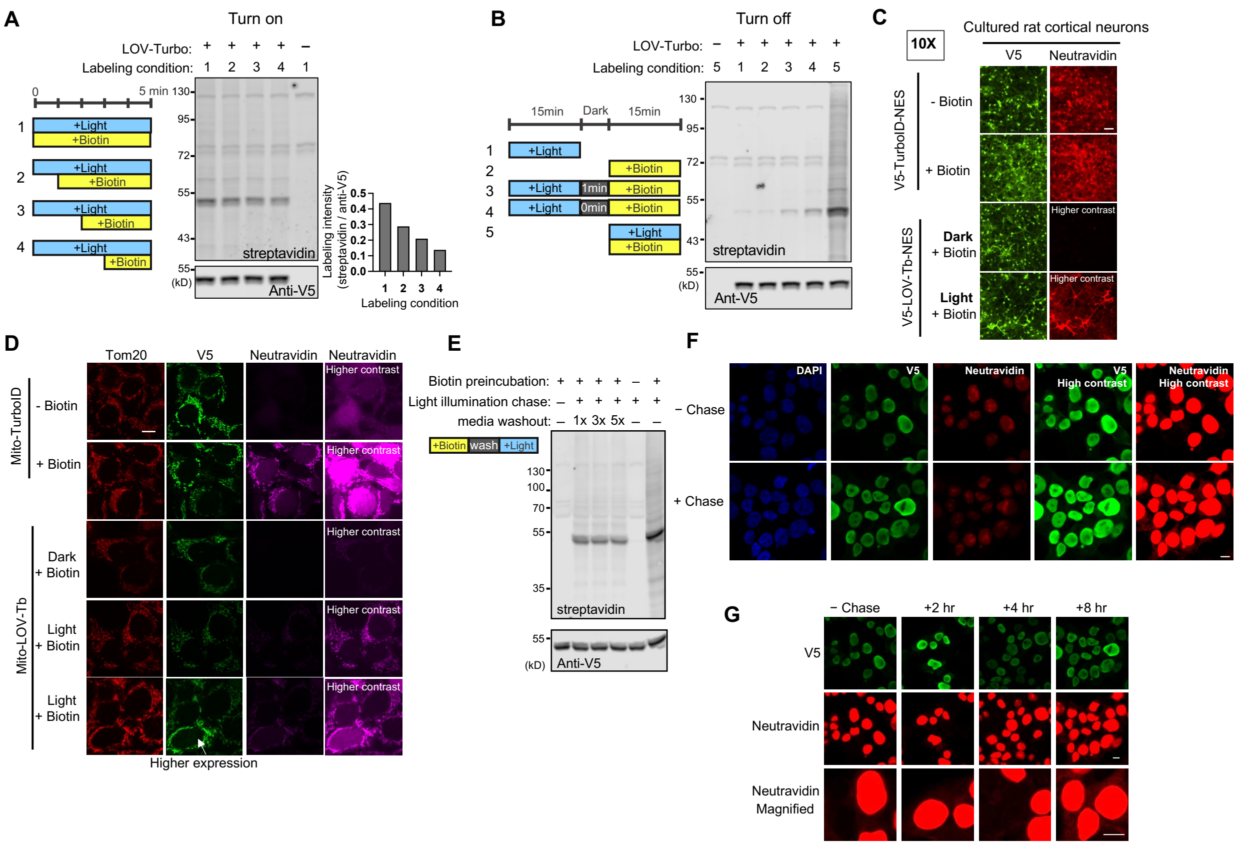


**Supplementary Fig. 3. Additional data related to LOV-Turbo characterization and applications.** (**A**) Kinetics of LOV-Turbo turn-on. Cytosolic LOV-Turbo was expressed in HEK 293T cells and labeling was performed as indicated. Labeling intensity drops as the biotin time window is shortened, indicating that LOV-Turbo turns on within a minute of exposure to blue light. (**B**) Kinetics of LOV-Turbo turn-off. Cytosolic LOV-Turbo was expressed in HEK 293T cells and labeling was performed as indicated. Light followed by biotin does not give significantly more labeling than biotin-only treatment, indicating that light removal turns off LOV-Turbo within a minute. (**C**) Same as **Fig. 3D**, but samples were analyzed by microscopy rather than streptavidin blotting. Scale bar, 100 µm. (**D**) Comparison of mitochondria-targeted TurboID and LOV-Turbo. Cells were labeled for 30 minutes, then fixed and stained. Scale bar, 10 µm. (**E**) Biotin washout is incomplete. HEK 293T cells expressing cytosolic LOV-Turbo were pretreated with biotin for 10 minutes and then lysed or washed with DPBS for the indicated number of times. After washing, cells were irradiated for 30 minutes in media lacking biotin. (**F**) Same as **Fig. 4C**, but without magnification. Scale bar, 10 µm. (**G**) Same as **Fig. 4C**, but with varying chase times. Scale bars, 10 µm.

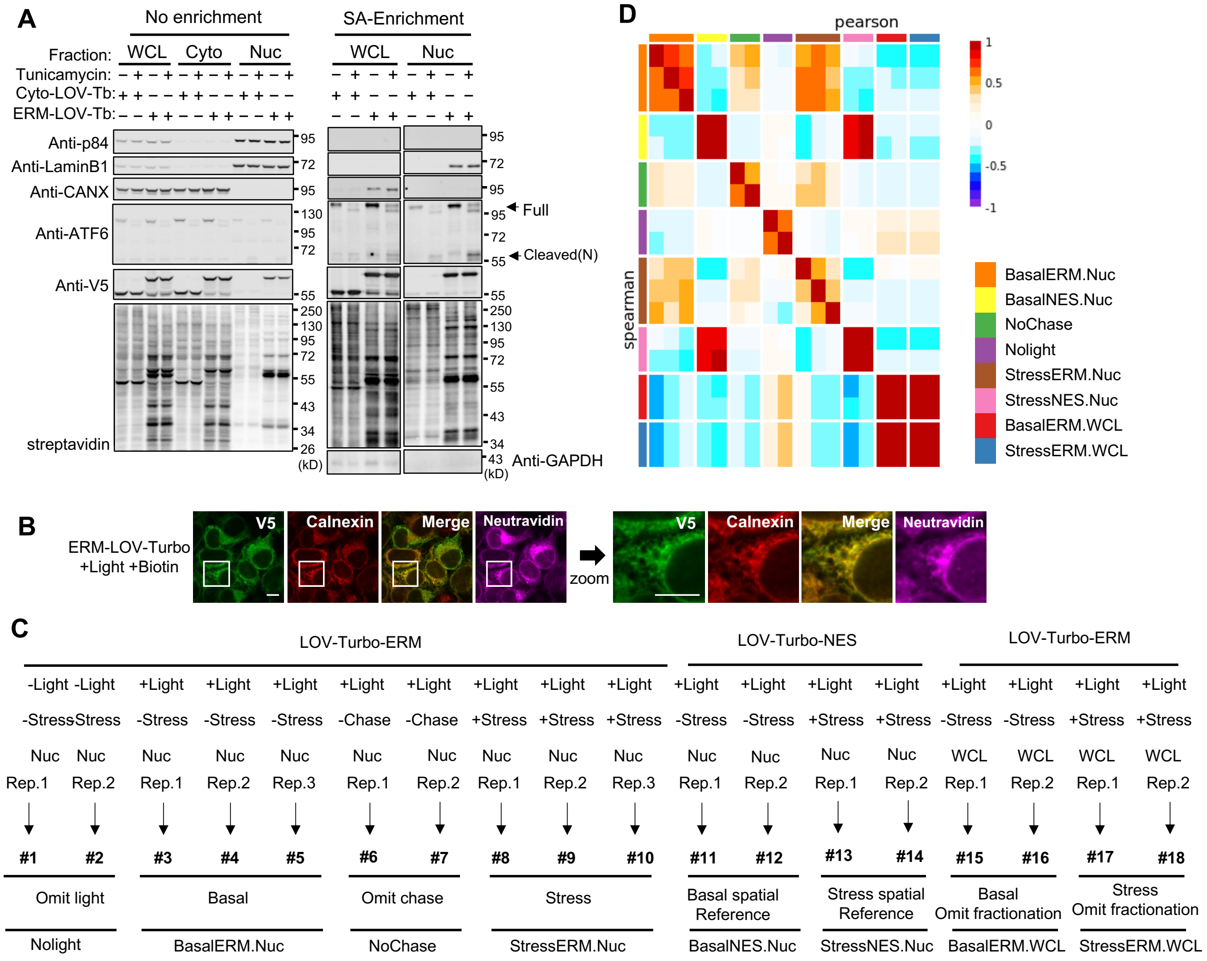
A
No enrichment
SA-Enrichment
Fraction: WCL Cyto Nuc
WCL Nuc
Tunicamycin:
Cyto-LOV-Tb:
ERM-LOV-Tb:
Anti-p84
Anti-LaminB1
Anti-CANX
Anti-ATF6
Anti-V5
streptavidin
Full
Cleaved(N)
Anti-GAPDH
(kD)
B
ERM-LOV-Turbo
+Light +Biotin
V5
Calnexin
Merge
Neutravidin
zoom
D
pearson
spearman
BasalERM.Nuc
BasalNES.Nuc
NoChase
Nolight
StressERM.Nuc
StressNES.Nuc
BasalERM.WCL
StressERM.WCL
C
LOV-Turbo-ERM
LOV-Turbo-NES
LOV-Turbo-ERM
-Light -Light +Light +Light +Light +Light +Light +Light +Light +Light +Light +Light +Light +Light +Light +Light +Light +Light
-Stress Stress -Stress -Stress -Stress -Chase -Chase +Stress +Stress +Stress -Stress -Stress +Stress +Stress -Stress -Stress +Stress +Stress
Nuc Nuc Nuc Nuc Nuc Nuc Nuc Nuc Nuc Nuc Nuc Nuc Nuc Nuc WCL WCL WCL WCL
Rep.1 Rep.2 Rep.1 Rep.2 Rep.3 Rep.1 Rep.2 Rep.1 Rep.2 Rep.3 Rep.1 Rep.2 Rep.1 Rep.2 Rep.1 Rep.2 Rep.1 Rep.2
#1 #2 #3 #4 #5 #6 #7 #8 #9 #10 #11 #12 #13 #14 #15 #16 #17 #18
Omit light
Basal
Omit chase
Stress
Basal spatial Reference
Stress spatial Reference
Basal Omit fractionation
Stress Omit fractionation
Nolight
BasalERM.Nuc
NoChase
StressERM.Nuc
BasalNES.Nuc
StressNES.Nuc
BasalERM.WCL
StressERM.WCL

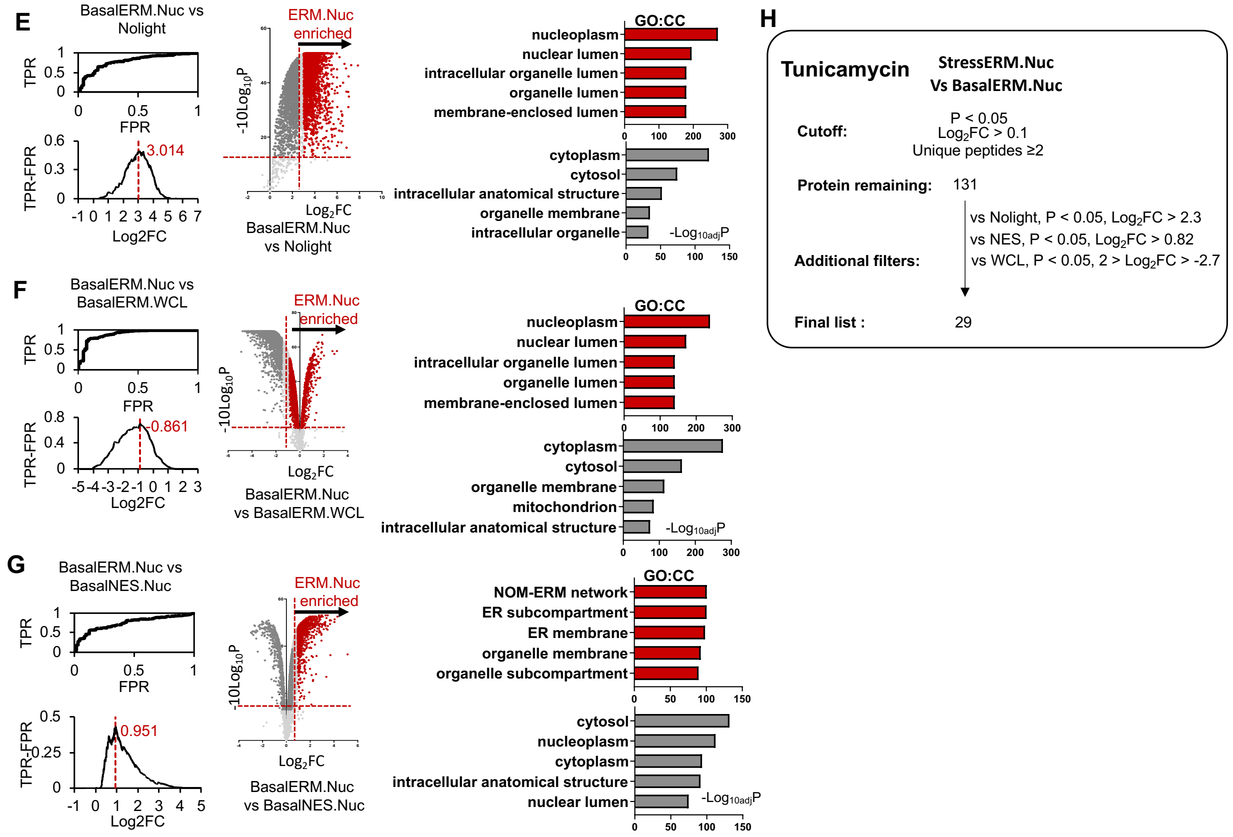


**Supplementary Fig. 4. Pulse-chase labeling with LOV-Turbo to discover proteins that traffick from ERM to nucleus.** (**A**) Western blot validation of labeling and nuclear fractionation procedure. Samples were prepared as in **Fig. 5A**. In the streptavidin (SA)-enriched material, we blotted for the true positive ERM-to-nucleus translocated protein ATF6, and found it cleaved in the nuclear fraction after tunicamycin treatment (arrows). Other antibodies used are anti-V5 (to detect LOV-Turbo-ERM), anti-p84 (to detect a nuclear matrix marker), anti-LaminB1 (a nuclear membrane marker), and anti-CANX (ER membrane marker). (**B**) ERM-LOV-Turbo is cleanly localized to the ER. Calnexin is an endogenous ER marker. Even though small portion of ERM-LOV-Turbo is cleaved and shows additional band in anti-V5 blotting shown in (A), confocal imaging shows clear ER targeting of ERM-LOV-Turbo. Scale bars, 10 µm. (**C**) Design of 18-plex TMT proteomic experiment to map ER membrane-to-nucleus translocation. +Stress samples were treated with 2 µg/mL tunicamycin. “Nuc” indicates that nuclear fractionation was performed. WCL, whole cell lysate. At bottom, sample names used in (D)-(H) and **Supplementary Table 2**. (**D**) Correlation between replicates. (**E-G**) Receiver operating characteristic (ROC) curves for various TMT ratios, selection of fold-change cut-offs (TPR-FPR plots), and volcano plots. At right, GOCC term enrichment analysis for filtered proteins (red in volcano plot) and excluded proteins (grey in volcano plot). In (E-F), TPs are nuclear proteins, and FPs are mitochondrial proteins. In (G), TPs are ER membrane proteins and FPs are mitochondrial proteins (see **Methods**). NOM, nuclear outer membrane. (**H**) Summary of filtering procedure used to generate final dataset of 29 stress-dependent ERM-to-nucleus translocated proteins.

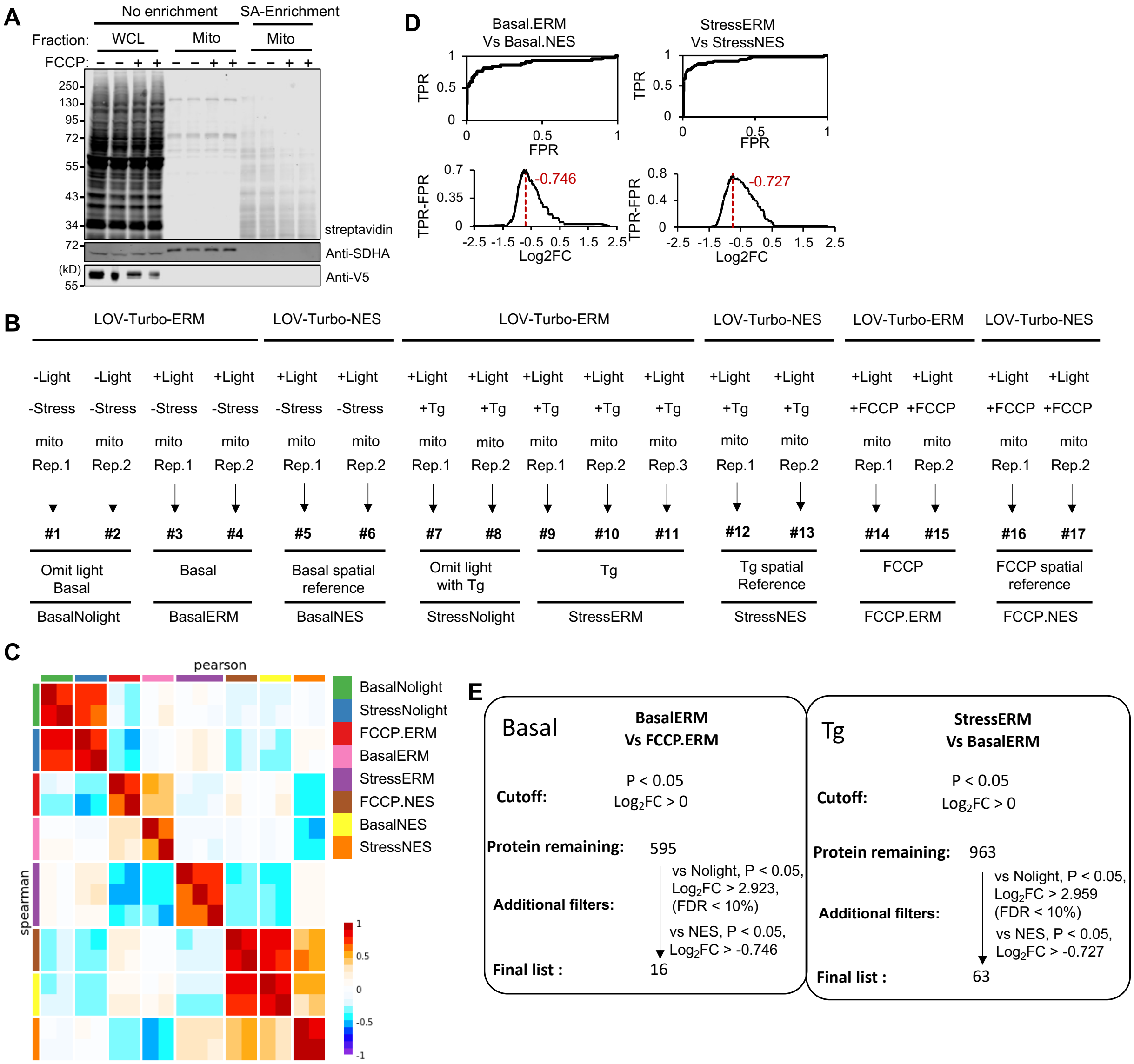


**Supplementary Fig. 5. Pulse-chase labeling with LOV-Turbo to discover proteins that traffick from ERM to mitochondria.** (**A**) Inhibition of mitochondrial protein import with FCCP. Samples were prepared as in **Fig. 5C**. Whole cell lysates (WCL), mitochondrial fractions, and streptavidin (SA)-enriched mitochondrial fractions were blotted with streptavidin-HRP, anti-V5 and anti-SDHA (an endogenous mitochondrial matrix protein marker). (**B**) Design of 17-plex TMT proteomic experiment to map ER membrane to mitochondria protein translocation. Tg, thapsigargin. At bottom, sample names used in (C)-(E) and **Supplementary Table 3**. (**C**) Correlation between replicates. (**D**) Receiver operating characteristic (ROC) curves for various TMT ratios (top), and selection of fold-change cut-offs (bottom). For TPR and FPR calculations, true positives were ER membrane proteins, and false positives were nuclear proteins (see **Methods**). (**E**) Filtering protocol for each dataset.

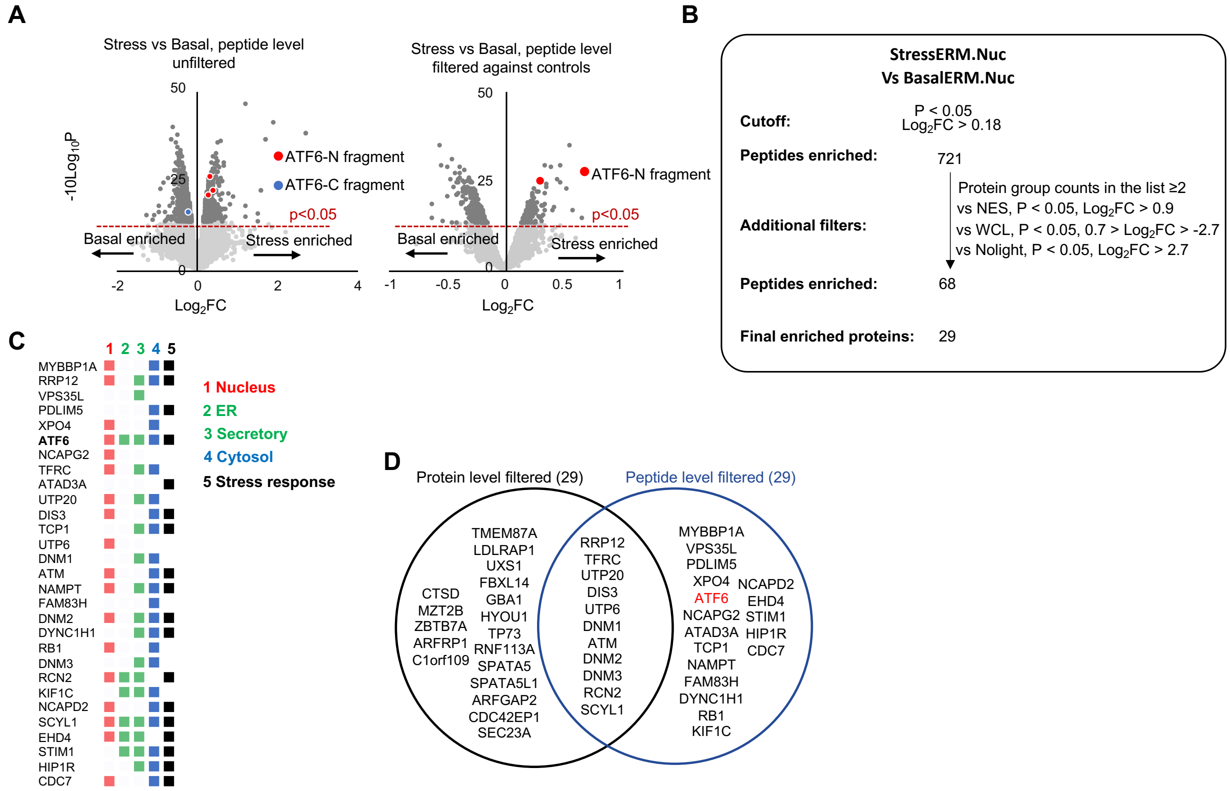


**Supplementary Fig. 6. Peptide level-analysis of ERM-to-nucleus proteomic data.** Related to **Supplementary Text 1**. (**A**) Volcano plots of peptide level data, unfiltered (left) or filtered against omit-light, cytosolic LOV-Turbo, and whole cell lysate control samples by peptide data (right). ATF6-derived peptides are highlighted in red (N-terminal fragment peptides, starting from amino acids 171, 202, or 217) or blue (C-terminal fragment-derived peptide starting from amino acid 405). (**B**) Filtering procedure for peptide data. First, stress condition peptide dataset was filtered against basal condition samples. This resulted in 721 unique peptide list. Then the list was grouped in protein groups, then unique peptides ≥2 protein groups were taken, and then further filtered against omit-light, cytosolic LOV-Turbo, and whole cell lysate control samples. (**C**) 29 proteins that exhibit increased trafficking from ERM to nucleus following tunicamycin, based on peptide level analysis as in (B). Proteins ranked by fold-change compared to basal condition. Colorings based on prior annotation (details in **Supplementary Table 4**). (**D**) Venn diagram showing overlap between protein and peptide level-filtered protein lists.

| | LOV domain | | | | | | | | | | | | | | | | | TurboID | | | | | | | | | | | | |
|---|---|---|---|---|---|---|---|---|---|---|---|---|---|---|---|---|---|---|---|---|---|---|---|---|---|---|---|---|---|---|
| | L8 | S33 | L37 | E39 | S41 | D71 | K89 | R100 | V106 | I110 | H121 | A123 | E125 | T135 | A136 | F137 | I139 | K2 | V6 | F51 | T52 | N68 | L73 | G74 | I147 | V156 | A166 | K168 | I231 | F286 |
| S10 | | | | | | | | | | | | | | | | | | | | | T52S | | | | I147T | | | | I231V | F286L |
| S16 | | | | | | | | | | | | | | | | | | | | | T52S | | | | I147T | | | | I231V | F286L |
| S17 | | | | | | | | | | | | | | | | | | | | | T52S | | | | I147T | | | | I231V | F286L |
| S18 | | | | | | | | | | | | | | | | | | | | | T52S | | | | I147T | | | | I231V | F286L |
| S1 | | | | | | | | | | | | | | | | | | | | | | | | | I147T | | | | | |
| S5 | | | | | | | | | | | | | | | | | | | | | | | | | I147T | | | | | |
| S3 | L6P | | | | | | | R98C | | | | A121T | | | | | I137T | | | | | | | | | | | | | F286L |
| S11 | L6P | | | | | | | R98C | | | | A121T | | | | | I137T | | | | | | | | | | | | | F286L |
| S19 | | | | | | | | | | | | | | | | F135S | | | | F51L | | | | | | | A166V | | | |
| S22 | | | | | | | | | | | | | | | | F135S | | | | F51L | | | | | | | A166V | | | |
| S2 | | | | | | | | | | | | | | | A134T | F135S | | | | | | | | | I147T | | | | | |
| S4 | | | | | | | | | | | | | | T133A | | | | | | | | | L73M | | | | | K168E | | |
| S6 | | S31P | | | | | | | | | | | | | | F135S | | | | | | | | | | | | | | |
| S7 | | | L35Q | | | | | | | | | | | | | | | | | | | | | | | | | | | |
| S8 | | | | | | | K87Q | | | | | | E123K | | | F135S | | K2E | | | | | | | I147T | | | | | |
| S9 | | | | | | | | | | | H119R | | | | | | | | | | | N68D | | | I147T | | | | | |
| S12 | | | | | | | | | | | | | | | | F135S | | | | | | | | | I147T | | | | | |
| S13 | | | | | | | | | | | | | | | | | | | | | | | L73Q | | | | | | I231V | |
| S14 | | | | E37G | | | | | V104I | | | | | | | | | | | | | | | | | | | | | |
| S15 | | | | | S39P | | | | | | | | | | | | | K2E | V6G | | | | | | | | | | | |
| S20 | | | | | | | | | | | | | | | | | | K2E | | | | | | | I147T | | | | | |
| S21 | | | | | | D69V | | | | I108T | | | | | | F135S | | | | | | | | | | | | | | |
| S23 | | | | | | | | | | | | | | | | | I137T | | | F51S | | | | G74E | | | | | | |
| S24 | | | | | | | | | | | | | | | | | I137T | | V6M | | | | | | | V156A | | | | |
| C1 | | | | | | | | | | | | | | | | | | | | | | | | | I147T | | | | I231V | |
| C2 | | | | | | | | | | | | | | | | | | | | | | N68D | | | I147T | | | | | |
| C3 | | | | | | | | | | | | | | | | | | | | | | | | | I147T | | | K168E | | |
| C4 | | | | | | | | | | | | | | | | | | | | F51L | | | | | I147T | | | | | |
| C5 | | | | | | | | | | | | | | | | | | | | | | | | | I147T | | A166V | | | |
| C6 | | | | | | | | | | | | | | | | | | | | | | | L73Q | | I147T | | | | I231V | |
| C7 | | | | | | | | | | | | | | | | | | | | | T52S | | | | I147T | | | | I231V | |
| C8 | | | | | | | | | | | | | | | | | | | | | | | | | I147T | | | | I231V | F286L |
| C9 | | | | | | | | | | | | | | | | | | | | | | | L73M | | I147T | | | | I231V | |
| C10 | | | | | | | | | | | | | | | | | | | | | T52S | | L73Q | | I147T | | A166V | | I231V | F286L |
| C11 | | | | | | | | | | | | | | | | | | | | | T52S | | L73Q | | | | A166V | | I231V | F286L |
| C12 | | | | | | | | | | | | | | | | | | | | | T52S | | L73Q | | I147T | | A166V | | I231V | |
| C13 | | | | | | | | | | | | | | | | | | | | | | | L73Q | | I147T | | A166V | | I231V | F286L |

C10 is selected as LOV-Turbo

**Supplementary Table 1.** LOV-Turbo variants enriched by directed evolution. Clones S1-24 are from round 7, and clones C1-13 are clones generated by manual combination of mutations. C10 is our final LOV-Turbo.

**Supplementary Table 2.** Proteomic data for ERM to nucleus pulse-chase experiment (**Fig. 5A**). **Tab 1**: 29 proteins that exhibit increased trafficking from ERM to nucleus following stress (tunicamycin) treatment. Proteins ranked by fold-change compared to basal (no stress) condition. Prior cytosol, nucleus, ER, secretory, and stress-related annotations listed. **Tab 2**: True positives (TPs) and false positives (FPs) used to check the specificity of the dataset as shown in **Supplementary Fig. 4E-G** and described in **Methods**. ERM TPs are ER-annotated proteins from GOCC; nuclear TPs are nuclear proteins used in Supplementary Table 3 (Tab 3) of Branon et al. Nature Biotechnology[14]. FPs are mitochondrial matrix proteins from GOCC. **Tab 3**: Unfiltered proteomic data showing fold-change and p-value for each pairwise comparison. **Tab 4**: Column definitions.

**Supplementary Table 3.** Proteomic data for ERM to mitochondria pulse-chase experiment (**Fig. 5C**). **Tab 1**: 16 proteins that traffick from ERM to mitochondria under basal condition. Proteins ranked by fold-change compared to FCCP condition. Prior integral membrane, secretory, MAM, mitochondria, ER, and cytosol annotations listed. **Tab 2**: 63 proteins that that exhibit increased trafficking from ERM to nucleus following stress (thapsigargin) treatment. Proteins ranked by fold-change compared to basal (no stress) condition. Prior integral membrane, secretory, MAM, mitochondria, ER, and cytosol annotations listed. **Tab 3**: True positives (TPs) and false positives (FPs) used to filter the proteomic data as shown in **Supplementary Fig. 5D-E** and described in **Methods**. ERM TPs are ER-annotated proteins from Supplementary Table 2 (Tab 1) of Branon et al. Nature Biotechnology[14]; nuclear TPs are nuclear proteins used in Supplementary Table 3 (Tab 3) of Branon et al. Nature Biotechnology[14]. FPs are mitochondrial matrix proteins from GOCC **Tab 4**: Unfiltered proteomic data showing fold-change and p-value for each pairwise comparison. **Tab 5**: Column definitions.

**Supplementary Table 4.** Proteomic data at peptide level for ERM to nucleus pulse-chase experiment (**Fig. 5A**). **Tab 1**: 29 proteins that exhibit increased trafficking from ERM to nucleus following stress (tunicamycin) treatment. Proteins ranked by fold-change compared to basal (no stress) condition. Prior cytosol, nucleus, ER, secretory, and stress-related annotations listed. **Tab 2**: Unfiltered proteomic data showing fold-change and p-value for each pairwise comparison. **Tab 3**: Column definitions.

#### Supplementary text 1. Further analysis of ERM-to-nucleus proteomic data.

ATF6 is a "true positive" protein that is known to translocate from the ERM to nucleus under stress[6, 7]. Though we detected it in our enriched material by Western blotting (**Supplementary Fig. 4A**), ATF6 was not in our final list of 29 stress-dependent ERM-to-nucleus translocated proteins. It may be absent from this list because it is cleaved in the process of translocation (only the N-terminal fragment arrives in the nucleus), resulting in fewer quantified peptides. Supporting this, volcano plot of peptide level data in **Supplementary Fig. 6A** shows N-terminal fragment peptide is enriched in stress condition. As other ERM-to-nucleus translocated proteins may undergo similar processing upon stress, we re-analyzed the MS data at the peptide, rather than protein level. To do this, as shown in **Supplementary Fig. 6B**, the 45956 detected unique peptides were filtered by p-value in addition to fold-change with respect to omit-stress (filter 1), cytosolic LOV-Turbo (filter 2), omit-light (filter 2) and omit nuclear fractionation (filter 2), in addition to protein group counts ≥ 2 within filter 1.

ATF6 is detected in the final list of 29 stress-responsive ERM-to-nucleus translocated proteins, due to strong enrichment of its N-terminal peptide (202-217 amino acids). Altogether, our list of 29 proteins (**Supplementary table 4**) contains 18 proteins with nuclear annotation, 17 with secretory pathway annotation, and 9 with both nuclear and secretory annotation (**Supplementary Fig. 6C**). 18 out of 29 proteins are known stress-responsive proteins. Four of our hits have previously been reported to translocate to the nucleus under stress (Exportin-4 XPO[8], Cyclic AMP-dependent transcription factor ATF-6 alpha ATF6[7], Serine-protein kinase ATM[9], and EH domain-containing protein 4 EHD4[10]).

Specific hits of interest include ATAD3A (ATPase family AAA domain-containing protein 3A), annotated as a mitochondrial inner membrane protein by GOCC. However, recent studies have assigned ATAD3A to mitochondria-ER contact sites[11] and we observed it to be enriched in our ERM-APEX2 dataset[12]. In addition, ATAD3A is responsive to ER stress[13] and truncated ATAD3A (1– 220 aa) localizes to the ER membrane[11]. We hypothesize that a C-terminal fragment of ATAD3A may be released by proteolysis and translocate to the nucleus under ER stress.

#### Antibodies used in this study

| Antibody | Source | Vendor | Catalog # | Dilution(s) |
|---|---|---|---|---|
| Anti-V5 | mouse | Invitrogen | R96025 | WB: 1:5000<br>IF: 1:3000 |
| Anti-SDHA | rabbit | Abcam | Ab137040 | WB: 1:1000 |
| Anti-TOMM20 | rabbit | Abcam | Ab186735 | WB: 1:3000<br>IF: 1:1000 |
| Anti-Hsp60 | rabbit | Cell Signaling | 4870 | WB: 1:3000 |
| Anti-p84 | rabbit | Abcam | Ab487 | WB: 1:3000 |
| Anti-calnexin | rabbit | Invitrogen | PA534754 | WB: 1:3000<br>IF: 1:1000 |
| Anti-ATF6 | rabbit | Proteintech | 24169-1-AP | WB: 1:3000 |
| Anti-LaminB1 | rabbit | Abcam | ab16048 | WB: 1:3000 |
| Anti-GAPDH-HRP | mouse | Santa Cruz Biotechnology | Sc47724 | WB: 1:2000 |
| Anti-6xHis | mouse | Thermo Fisher Scientific | MA1-21315 | WB: 1:1000 |
| Anti-Myc | chicken | Exalpha | ACMYC | Flow: 1:400<br>WB: 1:3000 |
| Anti-mouse-HRP | goat | BioRad | 1706516 | WB: 1:3000 |
| Anti-rabbit-HRP | goat | BioRad | 1706515 | WB: 1:3000 |
| Streptavidin-HRP | - | Invitrogen | S911 | WB: 1:3000 |
| Streptavidin-PE | - | Jackson ImmunoResearch | 0.16110084 | Flow: 1:200 |
| Neutravidin-AlexaFluor647 | - | Thermo Fisher Scientific | A2666; A20006 | IF: 1:1000 |
| DAPI | - | Enzo Life Sciences | AP402-0010 | IF: 1 µg/mL final concentration |
| Anti-mouse-AlexaFluor488 | goat | Invitrogen | A11029 | IF: 1:1000 |
| Anti-mouse-AlexaFluor568 | goat | Invitrogen | A11031 | IF: 1:1000 |
| Anti-mouse-AlexaFluor647 | goat | Invitrogen | A21236 | IF: 1:1000 |
| Anti-rabbit-AlexaFluor488 | goat | Invitrogen | A11008 | IF: 1:1000 |
| Anti-rabbit-AlexaFluor568 | goat | Invitrogen | A11036 | IF: 1:1000 |
| Anti-rabbit-AlexaFluor647 | goat | Invitrogen | A21245 | IF: 1:1000 |
| Anti-chicken-AlexaFluor488 | goat | Invitrogen | A-11039 | Flow: 1:200 |
| Anti-chicken-AlexaFluor647 | goat | Invitrogen | A21449 | Flow: 1:200 |
| Anti-Mouse-IRDye 800 | goat | LI-COR | 92632210 | WB: 1:5000 |
| Anti-Mouse-IRDye 680 | goat | LI-COR | 92668070 | WB: 1:5000 |
| Anti-Rabbit-IRDye 800 | goat | LI-COR | 92632211 | WB: 1:5000 |
| Anti-Rabbit-IRDye 680 | goat | LI-COR | 92568071 | WB: 1:5000 |
| Anti-Chicken-IRDye 680 | goat | LICOR | 92668075 | WB: 1:5000 |
| Streptavidin-IRDye 800 | - | LI-COR | 92532230 | WB: 1:5000 |
| Streptavidin-IRDye 680 | - | LI-COR | 92568079 | WB: 1:5000 |

### Genetic constructs used in this study

| Name | Promoter | Details | Used in |
|---|---|---|---|
| pCTCON2_Aga2P-LOV-Turbo1-Myc | GAL1 | | Fig. 1F and Supplementary Fig. 1E for yeast display |
| pTREtight_V5-LOV-Turbo variants-NES | TRE-tight | S1: I147T<br>S4: T133A, L73M, K168E<br>S9: H119R, N68D, I147T<br>S10: T52S, I147T, I231V, F286L<br>S12: F135S, I147T<br>S13: L73Q, I231V<br>S20: K2E, I147T<br>C1: I147T, I231V<br>C2: N68D, I147T<br>C3: I147T, K168E<br>C4: F51L, I147T<br>C5: I147T, A166V<br>C6: L73Q, I147T, I231V<br>C7: T52S, I147T, I231V<br>C8: I147T, I231V, F286L<br>C9: L73M, I147T, I231V<br>C10 (LOV-Turbo): T52S, L73Q, I147T, A166V, I231V, F286L<br>C11: T52S, L73Q, A166V, I231V, F286L<br>C12: T52S, L73Q, I147T, A166V, I231V<br>C13: L73Q, I147T, A166V, I231V, F286L | Fig. 1G-H and Supplementary Fig. 1F-H for transient expression in HEK 293T cells |
| pTRE3G-mito matrix (COX4I1)-LOV-Turbo variants-V5 | TRE3G | COX4I1 mito targeting sequence (MTS):<br>LATRVFSLVGKRAISTSVCVRAH<br>Amino acid differences across the LOV domains are colored red to facilitate comprehension. Linkers are highlighted in gray.<br>f-hLOV1 (I25V):<br>EF**R**ATTLERIEK**S**FVITDPRLPDNP**V**IF**V**SDSFLQLTEYSREEILGRNCRFL<br>QGPETDRATVRKIRDAIDNQTEVTVQLINYTKSGKKFWN**V**FHLQPMR<br>D**Y**KGDVQYFIGVQLDGTE**RLHG**AAEREAV**C**L**V**KKTA**FQ**I**A**<br>f-hLOV1 (V14T):<br>EF**R**ATTLERIEK**SFT**ITDPRLPDNPIIF**V**SDSFLQLTEYSREEILGRNCRFLQ<br>GPETDRATVRKIRDAIDNQTEVTVQLINYTKSGKKFWN**V**FHLQPMRD**Y**<br>KGDVQYFIGVQLDGTE**RLHG**AAEREAV**C**L**V**KKTA**FQ**I**A**<br>eLOV (143aa):<br>GS**R**ATTLERIEK**S**FVITDPRLPDNPIIF**V**SDSFLQLTEYSREEILGRNCRFLQ<br>GPETDRATVRKIRDAIDNQTEVTVQLINYTKSGKKFWNLFHLQPMRDQ<br>KGDVQYFIGVQLDGTE**R**VRDAAEREAVML**V**KKTAEEIDEAAK<br>eLOV (139aa):<br>GS**R**ATTLERIEK**S**FVITDPRLPDNPIIF**V**SDSFLQLTEYSREEILGRNCRFLQ<br>GPETDRATVRKIRDAIDNQTEVTVQLINYTKSGKKFWNLFHLQPMRDQ<br>KGDVQYFIGVQLDGTE**R**VRDAAEREAVML**V**KKTAEEID<br>WT LOV (1): GS-AsLOV2 (404-540aa):<br>GSLATTLERIEKNFVITDPRLPDNPIIFASDSFLQLTEYSREEILGRNCRFLQ<br>GPETDRATVRKIRDAIDNQTEVTVQLINYTKSGKKFWNLFHLQPMRDQ<br>KGDVQYFIGVQLDGTEHVRDAAEREGVMLIKKTAENID<br>WT LOV (2): EF-AsLOV2 (404-540aa):<br>EFLATTLERIEKNFVITDPRLPDNPIIFASDSFLQLTEYSREEILGRNCRFLQ<br>GPETDRATVRKIRDAIDNQTEVTVQLINYTKSGKKFWNLFHLQPMRDQ<br>KGDVQYFIGVQLDGTEHVRDAAEREGVMLIKKTAENID<br>iLID (Truncated): | Supplementary Fig. 1I for stable expression in HEK 293T cells |

| | | | |
|---|---|---|---|
| | | EFLATTLERIEKNFVITDPRLPDNPIIFASDSFLQLTEYSREEILGRNCRFLQ<br>GPETDRATVRKIRDAIDNQTEVTVQLINYTKSGKKFWN**V**FHLQPMRD**Y**<br>KGDVQYFIGVQLDGTE**RLHG**AAEREAV**C**LIKKTA**FQ**I**A**<br>hLOV1:<br>EF**R**ATTLERIEK**S**FVITDPRLPDNPIIF**V**SDSFLQLTEYSREEILGRNCRFLQ<br>GPETDRATVRKIRDAIDNQTEVTVQLINYTKSGKKFWN**V**FHLQPMRD**Y**<br>KGDVQYFIGVQLDGTE**RLHG**AAEREAV**CLV**KKTA**FQIA** | |
| pTRE3G_V5-LOV-Turbo1-NES | TRE3G | NES: LQLPPLERLTLD | Supplementary Fig. 1C for stable expression in HEK 293T cells |
| pTRE3G_V5-LOV-Turbo-NES | TRE3G | NES: LQLPPLERLTLD | Fig. 2B-E, 3A, 4B and 4E, and Supplementary Fig. 2A-G, 3A-B, 3E, 4C-D, 4G and 5C-D for stable expression in HEK 293T cells |
| pTRE3G_V5-LOV-Turbo-NLS | TRE3G | NLS: SRADPKKKRKVDPKKKRKVDPKKKRKV | Fig. 2C-E and 4C, and Supplementary Fig. 3F-G for stable expression in HEK 293T cells |
| pTRE3G_OMM(AKAP1(TM)-mito linker-GS linker)-V5-LOV-Turbo | TRE3G | AKAP1 transmembrane domain (TM):<br>MAIQLRSLFPLALPGLLALLGWWWFFSRKK<br>mito linker: DLELKLRILQSTVPRARDPPVAT<br>GS linker: GSSGGGSSGS | Fig. 2C-D for stable expression in HEK 293T cells |
| pTRE3G- V5-LOV-Turbo- ERM (SEC61B) | TRE3G | SEC61B:<br>PGPTPSGTNVGSSGRSPSKAVAARAAGSTVRQRKNASCGTRSAGRTTS<br>AGTGGMWRFYTEDSPGLKVGPVPVLVMSLLFIASVFMLHIWGKYTRS | Fig. 2C-D and 5A-E, and Supplementary Fig. 4A-H, 5A-E and 6A-D for stable expression in HEK 293T cells |
| pTRE3G-mito matrix (COX4I1)-LOV-Turbo-V5 | TRE3G | COX4I1 MTS: LATRVFSLVGKRAISTSVCVRAH<br>Mito matrix (1) | Fig. 2C-E, and Supplementary Fig. 2D and 3D for stable expression in HEK 293T cells |
| pTRE3G-mito matrix (COX4I1)-LOV-Turbo-V5-NES | TRE3G | COX4I1 MTS: LATRVFSLVGKRAISTSVCVRAH<br>Mito matrix (2) | Fig. 2D for stable expression in HEK 293T cells |
| pTRE3G-mito matrix (3x COX8A)-LOV-Turbo-V5-NES | TRE3G | 3x COX8A MTS:<br>MSVLTPLLLRGLTGSARRLPVPRAKIHSLQFSVLTPLLLRGLTGSARRLPV<br>PRAKIHSLARATMSVLTPLLLRGLTGSARRLPVPRAKIHSL<br>NES: LQLPPLERLTLD<br>Mito matrix (3) | Fig. 2D for stable expression in HEK 293T cells |
| pTRE3G-V5-TurboID-NES | TRE3G | NES: LQLPPLERLTLD | Fig. 2E and Supplementary Fig. 1C and 2B for stable expression in HEK 293T cells |
| pTRE3G-V5-TurboID-NLS | TRE3G | NLS: SRADPKKKRKVDPKKKRKVDPKKKRKV | Fig. 2E for stable expression in HEK 293T cells |

| | | | |
|---|---|---|---|
| pTRE3G-mito matrix (COX4I1)-V5-TurboID | TRE3G | COX4I1 MTS: LATRVFSLVGKRAISTSVCVRAH | Fig. 2E for stable expression in HEK 293T cells |
| pRS415_TurboID-V5 | GAL1 | | Fig. 3C for expression in BY4741 yeast strain<br>Addgene #107167 |
| pRS415_miniTurbo-V5 | GAL1 | | Fig. 3C for expression in BY4741 yeast strain<br>Addgene #107168 |
| pRS415_LOV-Turbo-V5 | GAL1 | | Fig. 3C for expression in BY4741 yeast strain |
| pET21a-TurboID-His6 | T7 | | Fig. 3B for expression in *E. coli*<br>Addgene #107177 |
| pET21a-miniTurbo-His6 | T7 | | Fig. 3B for expression in *E. coli*<br>Addgene #107178 |
| pET21a-LOV-Turbo-His6 | T7 | | Fig. 3B for expression in *E. coli* |
| pTRE3G-V5-miniTurbo-NES | TRE3G | NES: LQLPPLERLTLD | Fig. 2E and Supplementary Fig. 2B for stable expression in HEK 293T cells |
| pTRE3G-V5-miniTurbo-NLS | TRE3G | NLS : SRADPKKKRKVDPKKKRKVDPKKKRKV | Fig. 2E for stable expression in HEK 293T cells |
| pTRE3G-mito matrix (COX4I1)-V5-miniTurbo | TRE3G | COX4I1 MTS: LATRVFSLVGKRAISTSVCVRAH | Fig. 2E for stable expression in HEK 293T cells |
| pAAV_V5-TurboID-NES | CAG | NES: LQLPPLERLTLD | Fig. 3D and Supplementary Fig. 3C for AAV-induced expression in neurons |
| pAAV_V5-LOV-Turbo-NES | CAG | NES: LQLPPLERLTLD | Fig. 3D-E and Supplementary Fig. 3C for AAV-induced expression in neurons |
| pAAV_mito-TurboID-V5 | Synapsin | COX4I1 MTS: LATRVFSLVGKRAISTSVCVRAH | Fig. 4A for AAV-induced expression in neurons |
| pAAV_mito-LOV-Turbo-V5 | Synapsin | COX4I1 MTS: LATRVFSLVGKRAISTSVCVRAH | Fig. 4A for AAV-induced |

| | | | |
|---|---|---|---|
| | | | expression in neurons |
| pTRE3G-V5-NanoLuc-LOV-Turbo-NES | TRE3G | | Fig. 4G for transient expression in HEK 293T cells |
| pAAV_CCR6-NanoLuc-V5 | CMV | | Fig. 4I for transient expression in HEK 293T cells |
| pCDNA3_Myc-Arrestin-LOV-Turbo | CMV | | Fig. 4I for transient expression in HEK 293T cells |
